# Integrin-independent Tie2 activation using de novo designed proteins

**DOI:** 10.64898/2026.05.01.718274

**Authors:** Clara McCurdy, Yan Ting Zhao, Saurav Kumar, Brian Coventry, Anne Pink, Yaoyao Fu, Phil Bohn, Siyu Zhu, Inna Goreshnik, Xinru Wang, Garrett Ruth, Rashmi Ravichandran, Julie Mathieu, Jonathan A Cooper, Deborah H Fuller, Ho Min Kim, Pipsa Saharinen, David Baker, Hannele Ruohola-Baker

## Abstract

The Angiopoietin–Tie2 pathway is a key regulator of vascular stability, but therapeutic exploitation has been limited by the poor developability of Angiopoietin-1 (Ang1), and there are key unresolved mechanistic questions. Ang1 binds both Tie2 and α5β1 integrin, and the role of the α5β1 interaction in signaling has been unclear. We used RFdiffusion to design a stable, high-affinity Tie2 minibinder that has no affinity for α5β1. The binder is a selective antagonist in monomeric form, and a potent agonist when assembled into an octavalent architecture (H8mb) which drives Tie2 clustering. H8mb signals as potently as Ang1, indicating that integrin engagement is not required for Tie2 activation. However, the duration of signaling is reduced, and internalization of H8mb-Tie2 complexes is more rapid, suggesting that integrin may function as a co-receptor for Ang1 that prolongs signaling by extending the lifetime of the receptor ligand complex on the cell surface. In a mouse model of acute respiratory distress syndrome (ARDS), H8mb markedly improved survival. These results demonstrate that de novo designed receptor binders can enable dissection of co-receptor control of signaling dynamics, and the more stable and manufacturable H8mb provides routes to more developable therapeutic candidates.

---

The angiopoietin–Tie2 pathway is a central regulator of vascular and lymphatic stability, controlling endothelial survival, remodeling, and barrier integrity (*1–3*). The endogenous ligands Angiopoietin-1 (Ang1) and Angiopoietin-2 (Ang2) bind Tie2 through nearly identical fibrinogen-like domains yet produce opposing outcomes: Ang1 activates Tie2 and stabilizes vessels, whereas Ang2 antagonizes Ang1 signaling and promotes vascular destabilization (*1*,*4*). Disruption of the endothelial barrier contributes to diverse pathologies including traumatic brain injury, diabetic vasculopathies, retinopathies, and sepsis (*5–8*). Although several Tie2 agonists have been explored for vascular stabilization—including Ang1 variants, peptides, and antibodies—most suffer from limited specificity, poor manufacturability, or dependence on endogenous ligand expression (*9–12*). A longstanding unresolved question in this pathway is the role of integrin signaling. The Ang F-domain appears to interact not only with Tie2 but also with the adhesion receptor α5β1 integrin (*13*). Fibronectin has been reported to potentiate Ang1–Tie2 signaling, suggesting that integrin engagement may modulate Tie2 activation; however, the structural basis and functional necessity of integrin participation remain unclear (*14*). Because natural ligands engage both receptors, it has been difficult to disentangle Tie2 signaling from integrin crosstalk. We reasoned that this problem could be addressed by designing a ligand that exclusively binds Tie2. Recognizing that Tie2 signals through higher-order clustering, we separated the design challenge into two components: generating a high-affinity minibinder to the Tie2 extracellular domain and engineering a multimeric scaffold to oligomerize the binder. Constraining the design to the Ang binding interface ensured competition with endogenous ligands. Here we report H8mb, a de novo designed protein that acts as a potent and specific Tie2 agonist, enabling mechanistic dissection of Tie2 signaling independent of integrin engagement.

## RESULTS

### De novo design of a Tie2 specific minibinder

We reasoned that a Tie2 agonist could be designed de novo by combining two components - a small protein minibinder to specifically bind the receptor and an oligomerization domain to drive receptor clustering (*15*,*16*). We focused our minibinder design to the N terminal ‘arrowhead’ ligand binding domain of Tie2 (fig. 1A), which is highly conserved between mouse and human and serves as the binding site for the native ligands Ang1 and Ang2.The site had resisted earlier design campaigns because of its highly polar character and convex topology.

**Figure 1:**
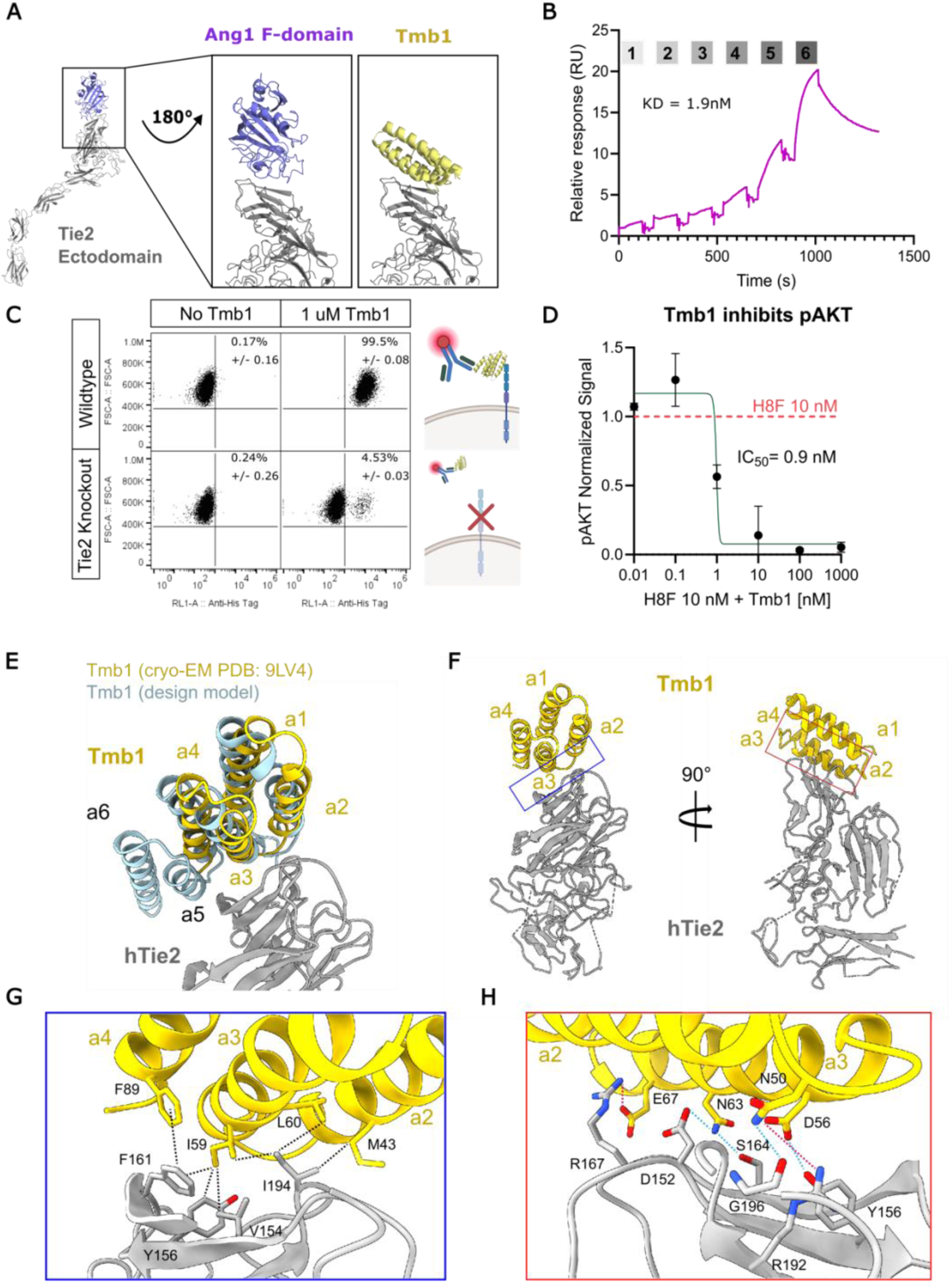
Design of a specific and high affinity Tie2 minibinder. A) Design strategy for the Tie2 minibinder showing that it was designed to compete with native F-domain binding from the ligands Ang1 and Ang2. B) SPR response trace for Tmb1 binding to human Tie2 ectodomain. Fitted *K*_D_ was calculated from 6 binder concentrations (0.016, 0.08, 0.4, 2, 10, and 50 [nM], labeled 1-6) using the Biacore Insight Evaluation software. C) Representative flow plots demonstrating that Tmb1 binds wild-type EA.hy926 cells, but does not bind to EA.hy926 cells with Tie2 knocked out. Percentages shown are the mean and standard deviation of 6 biological replicates from two independent experiments. D) Quantified western blot for pAKT in HUVEC cells. Blots were quantified using ImageJ and were normalized in-lane using S6 and then normalized to 10 nM H8F only. An example blot can be found in Fig. S5A. E) Cryo-EM structure (PDB ID: 9LV4) of the Tie2 minibinder Tmb1 (in yellow) overlaid with an Alphafold3 structure prediction model of Tmb1 (in blue) in complex with the ligand binding domain of human Tie2 receptor (in gray). F) Alternate views of the 4-helix cryo-EM structure of Tmb1. G-H) Close-up views of the interface residues resolved in the cryo-EM structure between Tmb1 and human Tie2.

Using a multi-step protocol combining RFDiffusion, ProteinMPNN, and AlphaFold2, we generated a library of designs targeting the LBD (see Methods). Yeast surface display identified 6 lead candidates which were then further optimized with site saturation mutagenesis and a second round of yeast display (fig. S1A-E). 96 combo-optimized designs were evaluated via surface plasmon resonance (SPR) to identify the highest affinity mutants of each of the 6 parents and the best mutants were tested via cell assays to identify our final candidate, Tmb1 (table S1). Ultimately, the RFDiffusion protocol was able to overcome previous limitations by creating concave binders with extensive shape complementarity to the convex target and large surface area thereby conferring enhanced affinity versus the previous attempts.

Tmb1 expressed efficiently in E. coli and eluted as a single monodisperse peak by size-exclusion chromatography (fig. S1F). Tmb1 has an estimated binding affinity of 1.6 nM for human Tie2 receptor (fig. 1B), which is stronger than the native ligand Ang1 (3nM, (4)). To assess specificity, we knocked down Tie2 in EA.hy926 endothelial cells using siRNA, incubated cells with Tmb1, washed excess binder, and detected cell-associated Tmb1 using an anti-His fluorophore. Flow cytometry analysis revealed that Tmb1 bound to wild type but not Tie2 knockout cells, indicating that Tmb1 does not bind other extracellular targets (fig. 1C). Residual binding to knockout cells coincides with the percentage of cells that still expressed Tie2 after knockout. In addition, Tmb1 also binds mouse Tie2. The extracellular domain of mouse and human Tie2 with an Fc tag were expressed in Expi293 cells. Tmb1 was co-expressed in the same cells with a 6x-His tag. When lysates were incubated on a ProteinA column and washed, Tmb1 eluted along with both types of Tie2, indicating that the minibinder bound to both mouse and human Tie2 (fig. S3). Finally, we tested whether Tmb1 competes with the Tie2 agonist H8F (*16*). Tmb1 successfully inhibited pAKT activation, demonstrating a potent IC_50_ of 0.9 nM. This high antagonistic ability suggests Tmb1 is binding to the intended F-domain/Tie2 interface, as originally designed (fig. 1A and 1D).

### Structural analysis of Tmb1-hTie2 binding

To elucidate the structural basis of Tie2 recognition by Tmb1, we determined the Cryo-electron microscopy (cryo-EM) structure of Tmb1 bound to the human Tie2 LBD. The hTie2 LBD and Tmb1 were co-expressed in HEK293F cells and purified by sequential Fc and His affinity chromatography, followed by SEC. Two cryo-EM maps were obtained from independent datasets. A lower-resolution 3.45 Å map revealed the overall architecture of Tmb1, with all six helices positioned as expected. A higher resolution 2.7 Å map enabled atomic modelling of helices α1–α4 of Tmb1, revealing side chain interactions with the hTie2 LBD (fig. 1E-H, fig. S2). Density corresponding to α5 in the AF3-predicted structure was only partially detectable in the 2.7 Å map and density for α6 was absent, suggesting that helices α5–α6 are not tightly integrated into the α1–α4 helical bundle.

The overall structure of the hTie2 LBD closely resembles its crystal structure (PDB ID: 2GY5, Cα RMSD of 1.13 Å; we did not build an atomic model of Tie2 in regions with low local resolution (fig. S2D). Tmb1’s relative position to hTie2 LBD is slightly deviated compared to the AF3-predicted Tie2 LBD/Tmb1 structure (fig. 1E). Ile59 of Tmb1 inserts into a hydrophobic cavity of Tie2, forming a stable hydrophobic network through interactions among residues on helices α2–α4 of Tmb1 (M43, I59, L60, and F89) and Tie2 residues (V154, Y156, F161, and I194) (fig. 1G–1H). These hydrophobic interactions were further stabilized by surrounding electrostatic interactions of N50, D56, N63, and E67 on Tmb1 α2–α3 with Tie2 residues (D152, Y156, S164, R167 and R192) (table S3).

Since the densities corresponding to helices α5–α6 of Tmb1 were barely detectable in the 2.7 Å map (fig. S2H), we wondered whether those two were still important for minibinder function. We hypothesized that they contribute to shielding a small hydrophobic patch centered around Phe161 on Tie2. To test this, we designed truncation variants lacking helices 5 and 6, as well as a disulfide-stabilized variant intended to rigidify the C-terminal region. We mutated up to 3 key non-interface surface residues to maintain solubility. All the mutants were expressed and purified and both truncations had a reduced ability to inhibit H8F, indicating reduced affinity (fig. S4A). Thus, despite their apparent flexibility, helices 5 and 6 are important for minibinder function consistent with the presence of all six helices in the lower-resolution 3.45 Å map (fig. S3H).

### Tmb1 is a tool to study Tie2 activity

Tie2 activation is strongly valency dependent: ligands presenting five or more F-domains induce robust Tie2/pAKT signaling and Ang1-like outputs (*16*). We therefore asked whether the high-affinity Tie2 minibinder (Tmb1) could substitute for the native Ang1 F-domain in multimeric scaffolds. To test this we conjugated Tmb1 to the octameric scaffold H8 using SpyTag and Spycatcher (*17*). The resulting molecule, designated H8-minibinder-spytag (H8mb^ST^), was successfully conjugated (fig. S4G) and, when applied to serum-starved HUVECs for a duration of 15 minutes, induced strong AKT phosphorylation with an EC₅₀ of 0.67 nM (fig. 2A).

**Figure 2:**
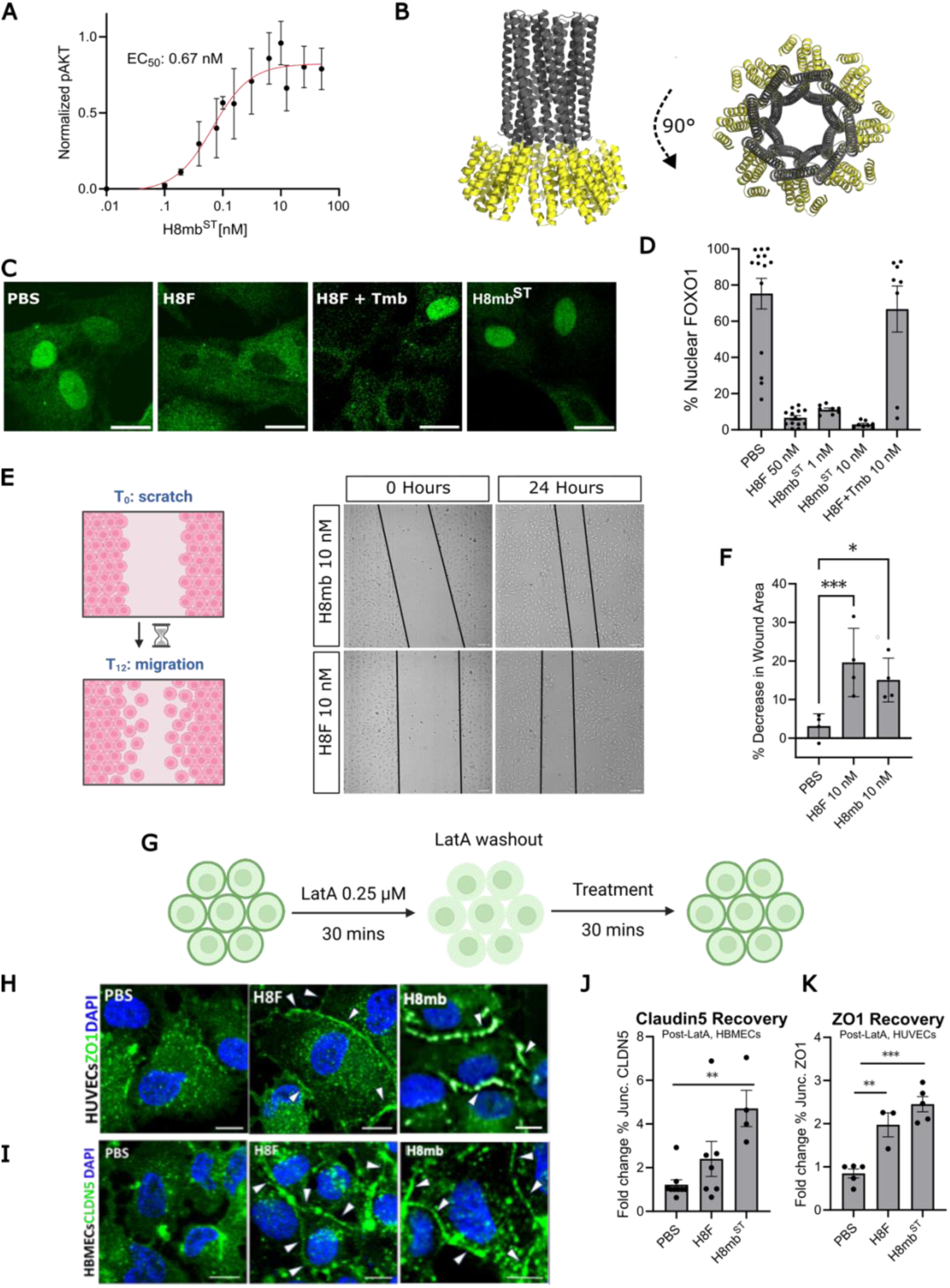
Multivalent H8mb activates Tie2. A) Quantified western blots for pAKT from serum-starved HUVEC lysate incubated with increasing concentrations of H8mb^ST^. The data was plotted and an EC50 curve derived from multiple experiments using Prism. Data are from at least 3 experiments and a representative blot can be found in S5C. B) Schematic of H8mb from two angles showing the presentation of 8 copies of Tmb1 (yellow) and the H8 scaffold (grey). C) Representative images of FOXO1 staining of HUVECs treated with (from left to right) PBS, 10 nM H8F, 10 nM H8F + 10 nM Tmb1 (Tmb), and 10 nM H8mb^ST^, scale bars are 50 µm. D) Quantification of HUVEC FOXO1 experiments presented as % of nuclei containing FOXO1. Each dot represents a biological replicate with at least 100 cells counted. E) Representation of the scratch assay in which a confluent monolayer of HUVECs are scratched and imaged at 0 hours and 12 hours post disruption. F) Representative images of scratch assay at 0 and 12 hours post scratch with cells incubated in either H8mb or H8F at 10 nM. G) Quantification of scratch assay results represented by the percentage by which the area of the scratch decreased. The area for each well was completed using the wound size healing tool in ImageJ (see methods). Each dot represents one biological replicate. H) Schematic demonstrating the LatA assay. Briefly, confluent HUVECs were treated with LatA for 30 minutes, washed, and then treated with PBS, 10 nM H8F, or 10 nM H8mb in low-glucose DMEM. I) Representative images of LatA experiment. HUVECs (top row) and HBMECs (bottom row) were fixed after the experiment and stained for ZO1 or CLDN5 and DAPI. The scale bar represents 10 µm. Quantification of LatA experiments for HUVECS (J) and HBMECs (K). Each dot represents a biological replicate with at least 100 cells counted. Briefly, the number of nuclei in each field of view were counted and each unique tight junction between cells was also counted. The ratio of nuclei and intact junctions were computed and the fold change between treatment groups is reported.

While H8mb^ST^ is a robust agonist, the SpyTag/SpyCatcher system is not desirable long-term due to the possibility of incomplete conjugation. Thus, we tested a variety of oligomer scaffolds by fusing Tmb1 to either the N or C termini of previously designed protein scaffolds with a GSGS linker between Tmb1 and the scaffold. Of the scaffolds tested and screened for pAKT activation (fig. S3B-F), the top candidate utilized the same H8 scaffold as above with the minibinder fused to the C-terminus and is called H8mb (fig. 2B).

A key downstream target of Tie2/pAKT signaling is the transcription factor FOXO1, which gets translocated out of the nucleus and degraded after stimulation (*3*). We therefore analyzed whether Tmb scaffolded at high valency has the capacity to induce FOXO1 nuclear exclusion. Serum-starved HUVECs were treated with H8mb^ST^ or H8F for 30 minutes and then fixed for antibody stains to identify the location of FOXO1 (fig. 2C-D). At concentrations as low as 1 nM, H8mb^ST^ almost completely depleted nuclear FOXO1, indicating robust activation of the pAKT/FOXO1 pathway. Conversely, co-treatment of HUVECS with H8F and monomeric Tmb1 restored FOXO1 to the nucleus across concentrations (fig. 2C-D), consistent with competitive binding of Tmb1 at the LBD interface and a higher apparent affinity than H8F.

### Oligomerized Tie2 mini binder promotes endothelial motility and tight junction repair

Having established the biochemical activity of H8mb in the Tie2 pathway, we next examined whether H8mb and H8F affect endothelial function. To test this, we set up a commonly used scratch wound assay to monitor cell migration over 12 hours (fig. 2E). We found that 10nM of H8mb and H8F both significantly increased cell migration to a similar degree (fig. 2F-G).

Secondly, to evaluate whether Tie2 activation modulates tight junction recovery, we disrupted endothelial tight junctions using Latrunculin-A (LatA) (*18*,*19*) in HUVECs and HBMECs. LatA inhibits actin polymerization which leads to the transient destabilization of tight junction protein complexes. Tight junctions are essential for maintaining cell-cell contacts and endothelial barriers, this method allows us to study their reformation as a proxy of Tie2 activity. Cells were treated with 0.25 uM of LatA for 15 minutes, washed, and then incubated with 10 nM of H8mb^ST^ or H8F for 30 minutes (fig. 2H-I). In HUVECs, both H8F and H8mbST increased junctional ZO-1 intensity more than two-fold over PBS control (fig. 2J). In HBMECs, H8mbST similarly enhanced recovery of claudin-5–positive junctions by approximately four-fold relative to control (fig. 2K). Together, these findings demonstrate that replacing the native F-domain with the de novo designed Tie2 minibinder Tmb1 recapitulates key biological outcomes of Tie2 activation, including pro-migratory signaling and rapid restoration of endothelial barrier components.

### F-domain binding promiscuity

As Tie2 agonists, H8F and H8mb trigger similar receptor activation. We sought to better understand their differences by studying the promiscuity of each receptor binding domain. To accomplish this we used two-dimensional (2D) protein sheets consisting of self-assembling A and B subunits that form micron-sized lattices, as previously described in (*16,20*). These sheets present multiple copies of a given cargo (Tmb1 or F-domain) and result in clusters of bound and recruited receptors that are large enough to detect via immunofluorescence (Fig 3A).

**Figure 3:**
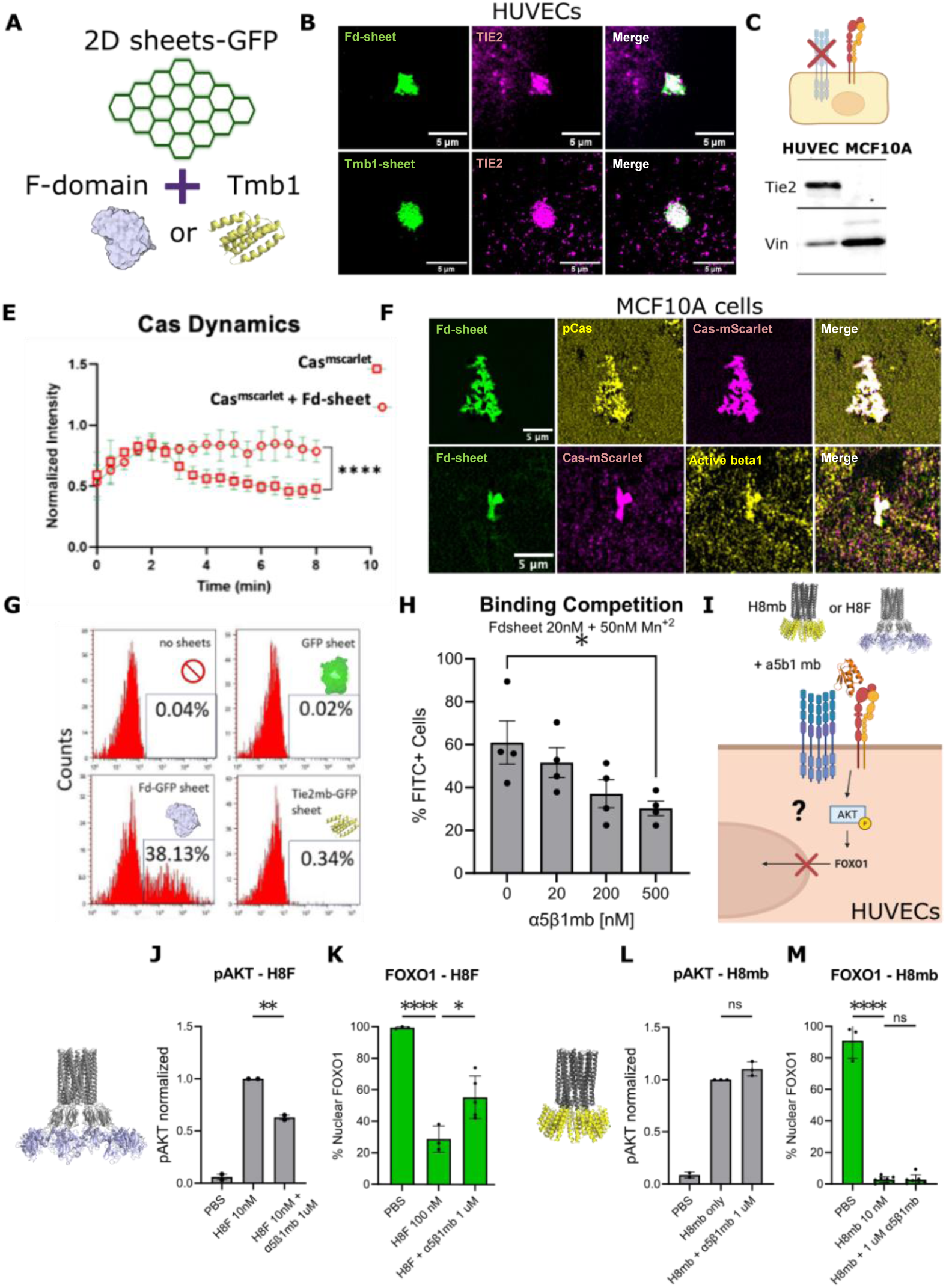
H8mb activates Tie2 without binding a5b1 integrin. A) Schematic representation of F-domain sheets which were made with GFP and either F-domain or Tmb1. B) Representative images of HUVECs with F-domain-sheets (top row) or Tmb1-sheets (bottom row) attached. Scale bars represent 5 µm. C) Schematic of MCF10A cells with representative western blot showing that MCF10A cells do not express Tie2. D) FACS analysis of MCF10A cells that were untreated or treated with 20 nM of either F-domain+GFP-sheets, Tmb1+GFP sheets, or GFP only-sheets for 15 minutes before PFA fixation. Representative histograms tracking FITC+ cells are shown. Buffers contained Mn^+2^ to potentiate integrin binding. E) Single channel and merged representative images of MCF10A cells that were treated with Fd-sheets for 30 minutes before staining and were imaged using a TIRF microscope. Cells were imaged to detect phosphoCas, total cas (Cas^mScarlet^) and active β1 integrin. Scale bars represent 10 um. F) Normalized Cas-mScarlet pixel intensity in a single location over 8 minutes on cells with no F-domain sheets (squares) and at the location an F-domain sheet is bound (circles). G) FACS analysis of Fd-sheet binding at 20 nM in the presence of α5β1 minibinder at 0, 20, 200, and 500 nM in MCF10A cells. H) Having investigated F-domain’s ability to recruit integrin in cells lacking Tie2 expression, the next question was whether integrin inhibition with the minibinder would affect downstream markers of Tie2 activation in cells that express both receptors. I) Quantified western blot for pAKT from HUVECs treated with PBS, H8F, or H8F + 1 uM α5β1. Each dot represents one biological replicate and is normalized to the total protein of the sample and then normalized to 10 nM H8F. J) Quantified FOXO1 staining of HUVECs treated with PBS, H8F 100 nM, H8F 100 nM + 1 uM α5β1, or 1 uM α5β1. Each dot represents one biological replicate with at least 100 cells counted. K) Quantified western blot for pAKT from HUVECs treated with PBS, H8mb, or H8mb + 1 uM α5β1. Each dot represents one biological replicate and is normalized to the total protein of the sample and then normalized to 10 nM H8mb. L) Quantified FOXO1 staining of HUVECs treated with PBS, H8mb 10 nM, or H8mb 10 nM + 1 uM α5β1. Each dot represents one biological replicate with at least 100 cells counted.

Tie2 interacts with integrin, specifically α5β1 integrin (*13*,*14,21*). The F-domain of Ang1 and Ang2 are thought to directly associate with Tie2 and α5β1, though the mechanism for integrin binding remains unclear. To confirm these interactions with F-domain and to test Tmb in this context, we conjugated either F-domain or Tmb1 to the protein sheets using SpyTag-SpyCatcher (*16*,*17*). Sheets were generated by conjugating GFP and F-domain (or Tmb1) to the A subunit at 1:4 ratio, respectively, so that GFP fluorescence would serve as a localization marker. The tagged A subunits were then combined with B subunits at a 1:1 ratio to produce the lattice. To confirm the binding of the sheets, HUVECs were incubated with either Tmb1- or Fd-sheets for 30 minutes. After washing, the cells were fixed and then stained with antibodies against Tie2. Both Fd- and Tmb1-sheets adhered to cells and clustered Tie2 as indicated by the co-localization of GFP and the Tie2 stains (fig. 3B).

Next, we probed the interaction of the native F-domain or Tmb1 with integrin. To do so, we moved to MCF10A cells which express integrins but lack Tie2 (fig. 3C). We first assessed whether F-domain binds integrin in MCF10A cells via flow cytometry. After gating on singlets and live cells, sheet positive cells were identified using the laser for FITC. Since Mn^2+^ ions enhance the affinity of α5β1 integrin for its native ligand fibronectin, which stabilizes the open confirmation (*22*,*23*), we reasoned that it might similarly promote F-domain binding and provide evidence that binding is integrin mediated. Indeed, the addition of 50 nM Mn^2+^ to the incubation buffer increased F-domain sheet attachment to around 30%, corresponding to an approximately 6-fold increase in the proportion of FITC+ cells relative to the sheet-only control, indicating F-domain binding to Integrin in MCF10 cells (fig. 3D & S5A). In contrast, Tmb1-sheets did not bind to MCF10A cells, further demonstrating that while Tmb1 binds Tie2 with high affinity, it does not bind Integrin (fig. 3D).

To determine whether F-domain is able to activate integrin when bound, we performed immunofluorescence to probe for focal adhesion markers co-localizing with the Fd-sheets. MCF10A cells stably expressing mScarlet–Cas (24) (Crk-associated substrate) were incubated with Fd-sheets and washed, fixed, and stained for active β1 integrin and phosphorylated Cas (pCas). Our data show that Fd-sheets colocalized with active β1 integrin, mScarlet–Cas, and pCas, indicating that clustered F-domain sheets induce integrin activation and focal adhesion assembly (fig. 3E). Furthermore, time-lapse microscopy analysis revealed that without an agonist present, Cas clusters spontaneously form and disassemble over minutes. We hypothesized that F-domain sheets bind and activate integrin leading to Cas clusters to persist for a longer duration. Indeed, in locations where Cas and Fd-sheets overlapped, Cas signal intensity stabilized for a significantly longer period (fig. 3F).

Lastly, confirming the specific role of α5β1 integrin in adherent cell lines through genetic knockout is difficult as cells tend to detach from the plate and die (unpublished). However, a recently developed monomeric and specific minibinder to α5β1 (α5β1mb) integrin that binds the RGD binding site provided a direct test of interaction specificity (*25*). We performed a competition assay with F-domain and α5β1mb to probe the model of F-domain-integrin binding. Co-incubating Fd-sheets with the α5β1 minibinder exhibited dose dependent inhibition of sheet binding on MCF10A cells using FACS analysis. 500 nM of binder decreased Fd sheet binding by half, demonstrating that the F-domain binding site is decreased when integrin is locked in the extended open conformation (fig. 3G).

### Activation of Tie2 by H8mb is independent of integrin

After confirming that F-domain binds α5β1 integrin but Tmb doesn’t bind in cells that lack Tie2, we next examined the effects in cells expressing both Tie2 and integrin to dissect whether integrin is needed for Tie2 activation, by using α5β1mb (fig. 3H). Serum-starved HUVECs were activated with 10 nM of H8F or H8mb, in the absence or presence of 1 uM α5β1mb to test pAKT/FOXO signaling. α5β1mb was able to repress H8F, but not H8mb induced pAKT/FOXO1 activity, further reiterating H8mb independence of α5β1 (fig. 3I-L)

### Tie2 activation by H8mb and H8F differs in duration of signal

Many receptor tyrosine kinases have co-factors or co-receptors that stabilize interactions with ligands and prolong signaling events (*26*,*27*). We investigated whether α5β1 integrin is needed to sustain Tie2 activation by analyzing the kinetics of AKT phosphorylation after H8mb or H8F administration. Using the same protocol as described above, HUVECs were stimulated up to 60 minutes (10 nM H8F or H8mb). Between 15 and 30 minutes, H8mb and H8F increased pAKT to a similar degree, but by 40-50 minutes, H8F exhibited significantly higher pAKT levels as compared to H8mb (fig. 4A-B), suggesting that H8F signal sustains longer than H8mb signal.

**Figure 4:**
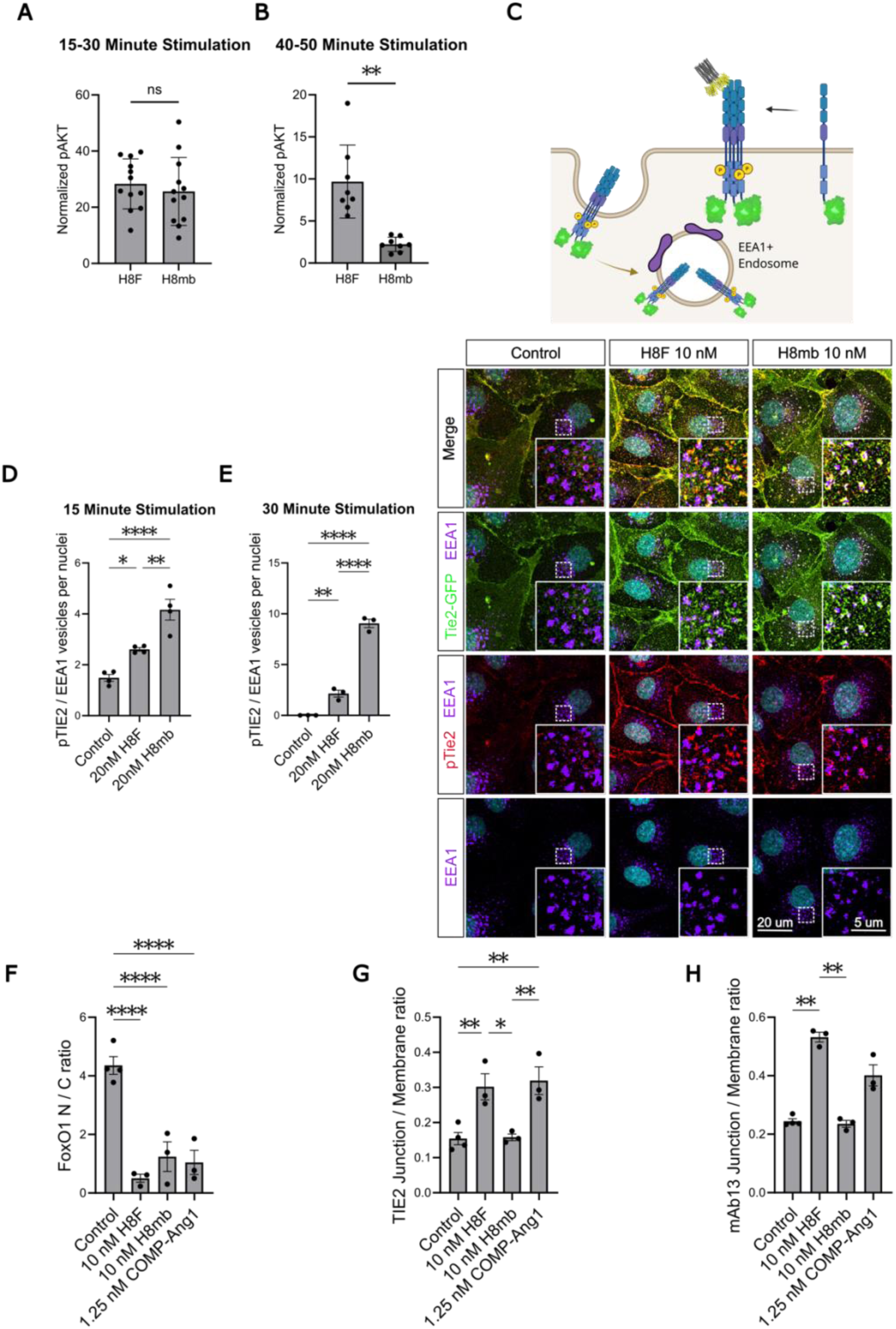
H8mb and H8F exhibit different signaling kinetics. A-B) Quantified western blot analysis of pAKT between 15-30 minutes and 40-50 minutes in HUVECs. Each data point represents one biological replicate and is normalized by per lane total protein. Significance tested with one way ANOVA. C) Representative images of serum-starved HUVECs that express Tie2-GFP and were treated with H8F and H8mb at 20 nM for 15 minutes. Cells were fixed and stained for DAPI, EEA1, and pTie2 to analyze the colocalization of EEA1 and Tie2/pTie2. D-E) Quantification of pTie2 and EEA1 colocalization in biological repeats for the 15 minute timepoint is found in C and for the 30 minute timepoint is found in D. F–H) Quantitative analysis of immunofluorescence experiments (in supplementary) analyzing Tie2 and inactive β1 integrin subcellular location in serum-starved HUVECs after 30 minutes of stimulation by 10 nM H8F or H8mb or 1.25 nM of another Tie2 agonist, Comp-Ang1. F) Quantitative analysis of FOXO1 reported as a ratio of nuclear to cytoplasmic FOXO1. G) Quantitative analysis of Tie2 signal from junctions vs non-junctions as identified by the binary mask identified by CDH5 staining (see methods). H) Quantitative analysis of inactive β1 integrin location reported as a ratio of junctional vs non-junctional signal. All statistical significances were assessed using ordinary one-way ANOVA followed by post hoc pairwise comparisons with Tukey’s multiple comparison test.

In order to further probe the kinetic differences between H8mb and H8F, we sought to track receptor internalization upon activation (fig.4C). Tie1 and other RTKs have been shown to relocalize to EEA1-containing early endosomes after stimulation (*28*,*29*) Serum-starved Tie2-GFP expressing HUVECs were treated with 20 nM of both H8mb and H8F for 15 or 30 minutes, fixed, and then stained for phosphoTie2 and EEA1 (fig. 4C). For both the 15 and 30 minute time points, H8mb treatment induced significantly higher intensity of pTie2 colocalizing in endosomes as compared to H8F, with the difference between the two appearing greater at 30 minutes (4E) than at 15 (4D). Total Tie2, as tracked by the location of Tie2-GFP, also was significantly increased in endosomes from cells treated with H8mb at both time points (fig. S7F-G). While the implications for Tie2 signaling for receptors found inside endosomes is unclear, taken together both the time course and the internalization experiments demonstrate an appreciable difference in the dynamics of the two ligands.

### H8mb and H8F differentially localize Tie2

Previously, Tie2 has been shown to colocalize with α5β1 integrin when activated by Ang 1 or Ang-like ligands (*14*,*16*). Since H8mb does not bind integrin, we wondered whether Tie2 or β1 integrin would still colocalize. Confluent, serum-starved HUVECs were treated with either 10 or 20 nM of H8F or H8mb or 1.25 or 2.5 nM of COMP-ang and then fixed and stained for CDH5, FOXO1, Tie2, and inactive β1 integrin. Representative images are shown in fig. S8-9 and multiple wells are quantified in figure 4F-H.

H8F, H8mb, and COMP-ang1 all excluded FOXO from the nucleus to a similar ratio and were significantly different from controls, indicating comparable activation (fig. 4F). To examine receptor subcellular localization, we used CDH5 staining to define junctional regions and create a binary mask and then computed the fraction of Tie2 and β1 integrin signal that overlapped with the mask. H8F and COMP-ang1 both similarly translocated Tie2 and β1 to junctions. H8mb meanwhile, did not significantly increase junctional Tie2 nor β1 integrin as compared to the control at either time point (fig. 4H & G and fig. S8 & S9), demonstrating that activation of Tie2 by H8mb induces less junctional distribution.

### H8mb prevents mortality in influenza infected mice

Damage to the lungs as a result of severe infection is a well-known cause of death associated with influenza and other infections. High inflammation in the lungs leads to a breakdown in endothelial barrier integrity and the subsequent microvascular leak can lead to respiratory failure and death (*30*). Tie2 has long been proposed as a potential therapeutic target for severe infection due to its role in endothelial barriers and an observed rise in inhibitory Ang2 in sepsis patients (*31*). Hence we sought to investigate whether H8mb confers therapeutic benefit in a severe influenza mouse model.

Female BALB(c) mice were inoculated with a 100x TCID50 dose of CA 2009 strain of H1N1 and treated intranasally with a vehicle control or with H8mb at 5 mg/kg starting one hour post-infection and continuing once per day for a total of 5 days (fig. 5A). Mice were monitored and their body weight measured every day for a week post inoculation. Mice in our higher dose treatment group survived at a significantly higher rate than the non-treatment group (fig. 5B). They also survived at significantly lower body weights. None of the control mice recovered their body weight if their weight dropped lower than 90% of their starting weight (fig. 5D) whereas several of the H8mb treated mice exhibited body weight dips as low as 75% of their starting weight but recovered body weight and survived the challenge (fig. 5C). These data indicate that H8mb provides a protective benefit in severe influenza, consistent with the idea that targeting Tie2 alone is sufficient to enhance barrier function in an inflammatory setting and that explicit integrin co-targeting may be dispensable for therapeutic efficacy.

**Figure 5:**
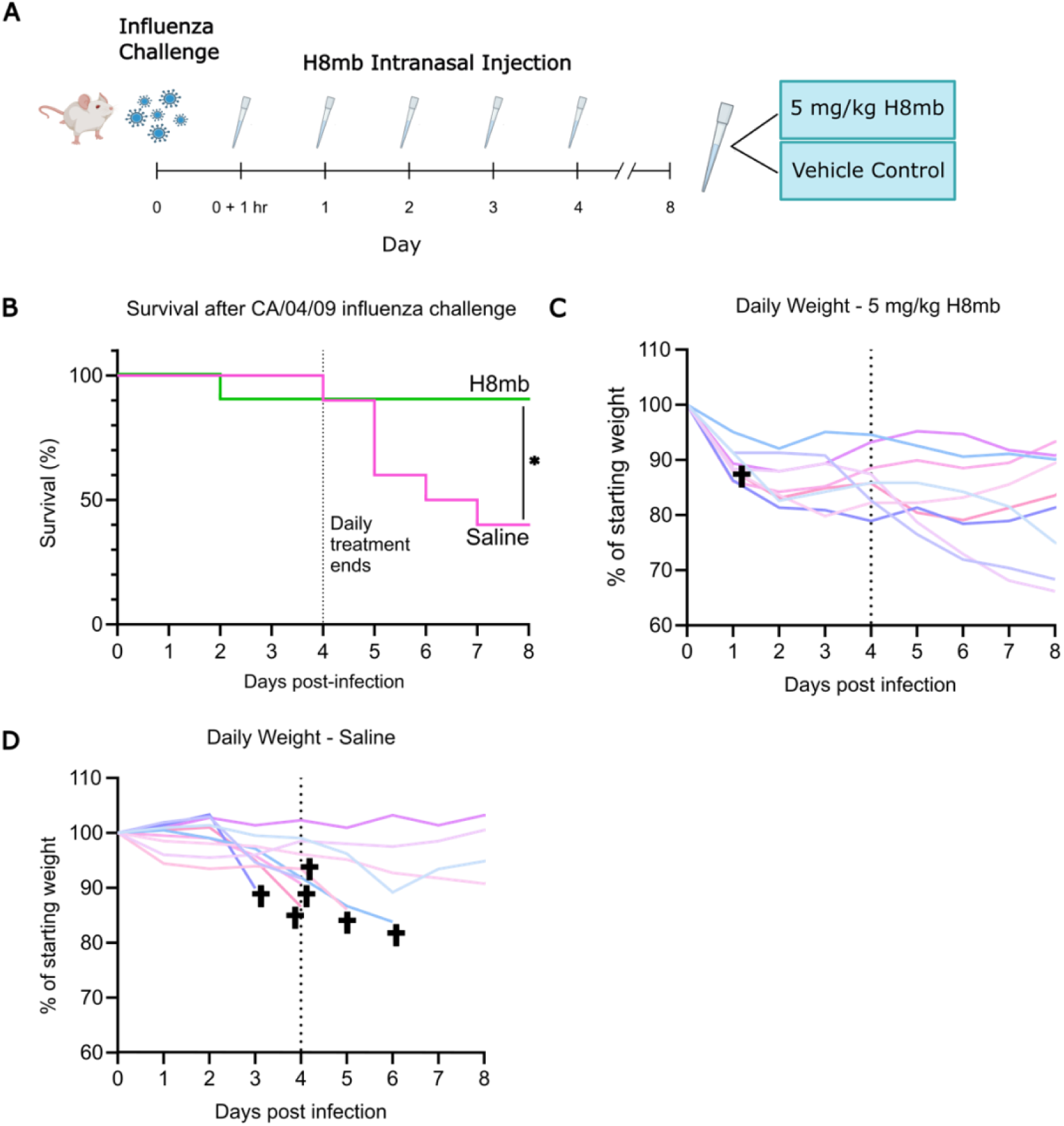
H8mb exhibits a protective influence over murine mortality in an ARDS model. A) Schematic representation of experimental lay out. Briefly, mice were infected with a 100x TCID50 dose of CA/04/09 influenza. Starting 1 hr post-infection and continuing 1x daily for 5 days, mice were dosed intranasally with a vehicle control or 2 mg/kg or 5 mg/kg of H8mb. B) Kaplan-Meier survival curve with significance testing showing that the higher dose group lessened mortality as compared to the saline control. C) Daily weight per mouse reported as % of starting weight for each mouse in the 5 mg/kg H8mb group (n=10). D) Daily weight per mouse reported as % of starting weight for each mouse in the saline control group (n=10).

## DISCUSSION

Our results indicate that engagement of α5β1 is not required for Tie2 activation, but instead likely enhances the stability and persistence of the signalling response. Activation of Tie2 by the multivalent, high affinity, and highly specific designed agonist H8mb robustly induced phopho-AKT and FOXO1 translocation out of the nucleus. In contrast, H8F signaling was attenuated by co-incubation with an α5β1-targeted minibinder, while H8mb remained unaffected. This highlights how Ang mimics, like H8F, depend on α5β1 co-engagement (via direct binding, conformational constraint, or other mechanisms) consistent with previous studies demonstrating that integrin augments Tie2 signaling (*13*,*14*). We propose that α5β1 integrin stabilizes cell-surface Tie2 clusters in order to sustain signaling and promote the physical reorganization of Tie2 to different subcellular locations. H8mb, which does not bind other surface proteins, exhibits shorter duration signaling and does not substantially translocate Tie2 to cell-cell contacts the way H8F does. H8mb also triggered rapid pTie2 endocytosis despite equivalent pAKT/FOXO1 signal, implying that α5β1 is a key modulator of trafficking that extends Tie2 signaling duration without being essential for signal initiation. Prior studies have implicated Integrin subunits in Tie2 trafficking; though conflicting evidence remains as to the mechanism of action and the specific subunits involved (11). Furthermore, despite these differences, both agonists restored junctional recovery after disassembly from LatA treatment, suggesting that junctional recovery is primarily driven by canonical pAKT/FOXO.

Our finding that H8mb treatment significantly improved survival in influenza-infected mice suggests that therapies targeting Tie2 are a promising new modality for treating sepsis. This view is consistent with prior work demonstrating that vascular leak and endothelial damage during infection are associated with an inflammatory response that can eventually culminate in death if left unchecked. By stabilizing the endothelial barrier through Tie2 activation, H8mb acts through a completely distinct mechanism from existing ARDS treatments which rely on supportive care (ventilation, oxygenation etc), antivirals to treat the underlying infection, or corticosteroids for their anti-inflammatory effects (32).

By binding selectively to Tie2 without interacting with α5β1, the designed Tie2 minibinder enables the dissection of signaling in a manner not possible for the native ligands. Collectively, our findings establish a platform for the modulation of Tie2 signaling and lay the groundwork for the development of advanced therapeutics directed toward this pathway.

## Materials and Methods

### Computational design of Tie2 binders

The Tie2 structure was obtained from PDB 2GY5 and trimmed down to just the native ligand site. RFDiffusion(33) was at first used to generate binders, but the 2-4 helical structures generated did not cover the hydrophobics on both sides of the beta-sheet which was an arbitrary goal of this design campaign. Instead, Secondary Structure and Block Adjacency Files (SS/ADJ files) were generated to encourage RFDiffusion to make 6 and 7 helical bundles wrapping around the beta sheet. The diffusion outputs were designed with MPNN and their structures predicted with AF2 initial guess (34). The AF2 structures were then further designed with MPNN and predicted again to improve metrics. While these generated structures met our shape goals, they didn’t meet the strict AF2 pae_interaction goals (pae_interaction < 8) we envisioned to generate binders to such a challenging target. Borrowing from the ideas in Cao 2022, the interfacial helices of the best 400 designs were extracted, new SS/ADJ files created to incorporate these helices into 6-7 helical bundles were created, and a new round of diffusion outputs were generated(35). The newly-grafted designs were then subjected to partial diffusion to increase interfacial diversity, designed with MPNN, predicted with AF2, designed with MPNN, and predicted with AF2 once more. This motif-graft protocol was repeated 2 additional times. The designs from all AF2 runs were pooled together and the 7,500 best designs by pae_interaction were ordered.

### Initial Tie2 Yeast Display Library

The 7,500 designs were reverse translated using DNAWorks and a 230bp 15,000 chip-synthesized oligonucleotide array was ordered from Agilent encoding these designs. The library of ∼400bp oligos was assembled in 12 pools using the method from Basanta et. al 2020 to yield the 7,500 unique oligos needed for testing(36). The 7,500 oligos were transformed into Saccharomyces cerevisiae EBY100 yeast and grown and sorted in a manner exactly the same as Cao et. al 2022(35). After analyzing the NGS, it was determined that since the designed lengths of the proteins spanned a huge range from 80aa-120aa, that something had gone terribly wrong and only designs of length <90aa were present in the library. A single binder was identified though (Tie2Bind_tw4000).

The 7,500 designs were then subjected to another round of yeast display. 1,500 of the designs showed enough data in the first library to be labeled confidently as non-binders and so a partial-diffusion library of 1,500 designs was used to potentially improve the single hit. This time, all designs were padded to 120aa before reverse translation by padding with random combinations of G,S,N,A,V, and Q (mostly favoring G and S) to ensure even PCR of all designs and to hopefully prevent mis-priming (by using random linkers rather than identically GGS (untested idea)). This time the yeast display showed all members of the library and 5 additional binders were discovered. The partial diffusion library of Tie2Bind_tw4000 did not improve the affinity of the design.

### Designed protein purification

Tie2 minibinders from the 96 well plate screen were purified as previously described(33). Briefly, minibinder genes were ordered as eBlocks (Integrated DNA Technologies) and inserted into customized vectors that added a 6xHis tag to the product using Golden Gate cloning. BL21 E. coli were transformed and grown overnight in small cultures in autoinduction media. Cells were collected via centrifugation and chemically lysed using BPER supplemented with 1% PMSF, lysozyme, and Benzonase. After centrifugation to clarify the lysate, proteins were purified using Immobilized Metal Affinity Chromatography, eluted off the columns in 200 uL, and separated by size via an AKTA FPLC (Cytiva). Proteins were run on a Superdex 75 5/150 Increase column (Cytiva 29148722). Candidate binders that aggregated significantly or did not express well were disregarded.

Once the top candidate binders were identified, designs were re-expressed in appropriately scaled up cultures and expressed using the low-endotoxin approach described previously(37).

Candidate oligomers were prepared in small scale the same way the candidate binders were with the exception of using a Superdex 200 Increase 5/150 column (Cytiva 28990945). The oligomer that yielded the best result (H8mb) was scaled up and produced by the General Protein Production Core at the Institute for Protein Design. H8mb was expressed in a low-endo protocol as previously described(37).

### Constructs for recombinant expression of Tie2/minibinder Tmb1 complex for cryo-EM

Human Tie2 ligand binding domain (residues M1-N452), followed by a thrombin cleavage site and an Fc-tag was cloned into the HindIII and XhoI sites of the pcDNA 3.1 vector (#V79020, Invitrogen), which includes its own signal sequence for protein secretion. The mouse Tie2 ligand binding domain was inserted in a similar manner. In addition, the tie2 minibinder candidates, followed by a 6×His tag, were cloned into the BamHI and XhoI sites of a pcDNA 3.1 vector containing the vascular endothelial growth factor receptor (VEGFR) signal sequence for protein secretion.

### Expression and purification of the hTie2/minibinder Tmb1 complex for cryo-EM

A total of 1 μg of plasmid DNA (hTie2-Fc tag: minibinder Tmb1-6×His at a 1:4 ratio) was transfected into 3.0×106 Expi 293F cells (#A14527, Thermo Fisher Scientific) using Expifectamine (#A14524, Thermo Fisher Scientific). The cells were cultured in Expi293 expression medium (#A14351, Thermo Fisher Scientific) at 37°C and 8% CO2 with shaking at 120 rpm (orbital shaker) for 4 days. After centrifugation to remove the cells, the supernatant was loaded onto ProATM rProtein A Agarose Resin (#1010100, Amicogen). The resin was washed with 10 column volumes of wash buffer (20 mM Tris-HCl pH 8.0, 200 mM NaCl), and the bound hTie2-Fc/minibinder Tmb1-6×His were incubated with thrombin (1% [v/v] in wash buffer) at 4°C for 12 h to remove the C-terminal Fc tag. The eluted hTie2/minibinder 1102-6×His complexes were then applied to Ni-NTA Agarose (# 30210, Qiagen). After washing with 10 column volumes of wash buffer (20 mM Tris-HCl pH 8.0, 200 mM NaCl; 20mM Imidazole) to remove the non-specific bound impurities, the complexes were eluted in buffer containing 20 mM Tris-HCl pH 8.0, 200 mM NaCl; 250mM Imidazole. This process was repeated for the bound mouseTie2-Fc/minibinder complexes. The eluted hTie2/minibinder Tmb1 complexes were concentrated to 1 mg/mL using an Amicon Ultra centrifugal filter (#UFC8030, Millipore) and further purified by SEC on a Superdex 200 Increase 10/300 GL column (#GE28-9909-44, Cytiva) equilibrated in 20 mM Tris-HCl pH 8.0, 200 mM NaCl. The peak fraction was used for cryo-EM analyses without additional concentration.

### Cryo-EM sample preparation and data collection

Quantifoil R1.2/1.3 Cu #300 mesh grids (Quantifoil Micro Tools) were glow discharged using a PELCO easiGlow Glow Discharge Cleaning system (Ted Pella) for 90 s at 15 mA. Approximately 3 ul of purified hTie2/minibinder Tmb1 complex was applied to the grids under 100% humidity at 4°C. After a 3 s blot, the grid was rapidly vitrified by plunging them into liquid ethane using a Vitrobot MkIV (Thermo Fisher Scientific). Micrographs were acquired on a TFS Krios G4 TEM operated at 300 keV with a K3 direct electron detector (Gatan) at the Institute for Basic Science, using 20 eV slit on a GIF quantum energy filter. EPU software was used for automated data collection in single-electron counting and correlated double sampling (CDS) modes. Detailed image acquisition parameters for the hTie2/minibinder Tmb1 complex are summarized in Supplementary Table 1.

### Image processing, model building, and refinement

Detailed image processing workflow and statistics are summarized in Supplementary Fig. 4. Raw movies were motion-corrected using MotionCorr2 and the contrast transfer function (CTF) parameters were estimated with CTFFIND4(38,39). All further processing was carried out in cryoSPARC v.4.2.(40). Initially, particles were picked with a blob picker, binned four times, and subjected to 2D classification, ab initio modeling, and heterogeneous refinement for data cleanup. Final particles from 3D classes with clear secondary structural features were re-extracted at the original pixel size. Non-uniform refinement(41), local motion correction(42), and CTF refinement(43) improved particle alignment and map quality, yielding final maps at approximately 3.0 Å resolution (with a tight mask). The mask-corrected Fourier shell correlation (FSC) curves were calculated in cryoSPARC, and reported resolutions were based on the gold-standard FSC = 0.143 criterion(44). Local resolutions were estimated by Blocres(45).

Model building for the hTie2/minibinder Tmb1 complex began by docking the crystal structure of the hTie2 ligand binding domain (PDB ID: 2GY5)(46) into the post-processed cryo-EM map generated by DeepEMhancer(47) in cryoSPARC. The Cɑ chain and side chains of the minibinder were manually built into the density map using Coot (v.0.9.4.1)(48), with guidance from its AlphaFold3-predicted structure. The resulting models were manually adjusted in Coot and further refined against the map using the real space refinement in the Phenix package(49). The refinement statistics from Phenix validation are summarized in Supplementary Table 1.

### H8mb Conjugation

Tie2 minibinder 1102 was expressed with a C terminal SpyTag003(50). After being expressed and purified as described above, the minibinder was incubated at a 1.2:1 ratio of minibinder to H8 oligomer subunit that was tagged with the SpyCatcher sequence at the C terminal. Incubation took place overnight at 4C on nutation. The H8F oligomer was prepared as described above and as was previously published in Zhao 2021 (16).

### Cell surface binding specificity analysis by flow cytometry

EA.hy926 adherent cells (ATCC CRL-2922) were cultured in T75 at 37°C in a 5% CO2 atmosphere in Dulbecco’s minimum essential medium (DMEM; ThermoFisher11885084) supplemented with 10% heat-inactivated Hyclone fetal clone II serum and 1% penicillin/streptomycin. Tie-2 knockout cells were generated using sgRNA and spCas9 obtained from Gene Knockout Kit v2 (Synthego). Cells were nucleofected with a Lonza 4D-Nucleofector Unit using the Lonza SF Cell Line 4D-Nucleofector™ X Kit according to the manufacturer’s protocol. To perform the specificity assay, cells were trypsinized, quenched with media, and transferred to a 96-well U-bottom plate. Cells were washed with DPBS supplemented with 0.2% BSA (FACS buffer) and incubated with varying concentrations of minibinder protein on ice for 30 min. Cells were washed with FACS buffer and incubated on ice with solutions of either CoraLite® Plus 647-conjugated his-tag monoclonal antibody (ProteinTech CL647-66005) or human Tie-2 Alexa Fluor® 647-conjugated antibody (R&D Systems FAB3131R). Following a 30 min incubation, cells were washed and resuspended in a Sytox Green/FACS buffer solution. Flow cytometry was performed using Attune CytPix Flow Cytometer and FlowJo software was used to gate on single cells and live cells for analysis.

### Cell culture

Human Umbilical Vein Endothelial Cells (HUVECs) were obtained from Lonza (C2519AS). Cells were grown on 0.1% gelatin coated plates in the Lonza EGM-2 Microvascular BulletKit® Media (CC-3202) which was prepared as directed. Cells were first expanded to passage 4 and then cryogenically preserved until needed. Unless indicated otherwise, all experiments with HUVECs were conducted between passages 7-8.

MCF10A cells were cultured in media described previously54,56; briefly, the media consisted of DMEM/F12 (Gibco, 11330032), 5% horse serum (Gibco, 16050130), 20ng/ml EGF (Sigma-Aldrich, SRP3027), 0.5mg/ml hydrocortisone (Sigma-Aldrich, H4001), 100ng/ml cholera toxin (Millipore, C8052), 10ug/ml insulin (Sigma-Aldrich,11070-73-8) and 1% penicillin–streptomycin (Gibco, 15140122). MCF10A cells were starved in the same media without EGF and containing 2% horse serum (assay media) for 16 hours before experiments.

### Total Protein Isolation

HUVECs were grown to at least 80% confluence in 12-well plates. They were then rinsed twice with PBS and starved in Low-glucose DMEM for 16-20 hours. After treatment with study molecules for the indicated duration, cells were washed with PBS once and were lysed with 65 μl of lysis buffer containing 20 mM Tris–HCl (Sigma-Aldrich, 1185-53-1) (pH 7.5), 150 mM NaCl, 15% glycerol (Sigma-Aldrich, G5516), 1% triton (Sigma-Aldrich, 9002-93-1), 3% SDS (Sigma-Aldrich, 151-21-3), 25 mM b-glycerophosphate (Sigma-Aldrich, 50020-100G), 50 mM NaF (Sigma-Aldrich, 7681-49-4), 10 mM sodium pyrophosphate (Sigma-Aldrich, 13472-36-1), 0.5% orthovanadate (Sigma-Aldrich, 13721-39-6), 1% PMSF (Roche Life Sciences, 329-98-6), 25 U benzonase nuclease (EMD, 70664-10KUN), and phosphatase inhibitor cocktail 2 (Sigma-Aldrich, P5726). Lysates were then frozen for further analysis.

### Western Blotting

Cell lysates were analyzed using one of two methods. Fig. 1I, 2A: Lysates were diluted 1:4 in 4x Laemmli Buffer (Bio-Rad #1610747) and then boiled at 95 °C for 10 minutes. Samples were run on a 4-10% SDS-PAGE gel for 30 minutes at 250 volts and transferred to a nitrocellulose membrane for 12 minutes at 2.5 volts using a semi-dry transfer machine (Bio-Rad). After transfer, the membranes were blocked in 5% bovine serum albumin for 1 hour. The membranes were then incubated overnight with anti-pAKT (1:1000) and/or a housekeeping gene, either S6 or β-actin (1:10,000). The next day, membranes were washed 3 times with TBST and incubated with the appropriate secondary antibody. Membranes were developed using chemiluminescent HRP substrate as directed (MilliporeSigma, WBKLS0050) and imaged with a ChemiDoc Imager (Bio-Rad). Blots were quantified using the ImageJ (FIJI) band intensity analyzer. Target protein intensities were normalized to the corresponding housekeeping gene from the same sample.

Some lysates were analyzed using a Jess automated western blot machine. Cells were lysed in either 65 or 30 uL of Mammalian Protein Extraction Reagent (ThermoScientific 78503) with 1% PMSF (Roche Life Sciences 329-98-6) and phosphatase inhibitor cocktail 2 (Sigma-Aldrich P5726) added in. Samples were run undiluted using the reagents provided as part of the 12-230 kDa separation module (Bio-techne SM-W001) and the anti-rabbit detection module (Bio-techne DM-001). Target proteins were probed using anti-pAKT antibody at 1:30 dilution. Target proteins were normalized using either β-actin (1:250 dilution) or using the Total Protein Normalization Module (Bio-techne DM-PN02). The machine was run with the appropriate default settings.

### Scratch Assay

The scratch assay was performed exactly as described in Zhao et al 2021. Briefly, HUVECs were plated in 12-well plates and grown to confluency. Once confluent, the cells were washed with PBS, and the media was changed into low-glucose DMEM with 2% fetal bovine serum. A scratch was made with a 200uL pipette tip, and phase-contrast images were taken in the same spots, two per well, at 0 hours and 12 hours. The area of the scratch was quantified in ImageJ (FIJI) using the wound healing size tool as described in Suarez-Arnedo et al (51).

### TIRF live-cell and immunofluorescence microscopy

Dual-color live cell imaging was performed immediately after plating cells till 30 min of time period. The images were recorded using TIRF microscope (Nikon Ti, 100x/1.49 CFI Apo TIRF oil immersion objective) equipped with Perfect Focus, motorized x-y stage, fast piezo z stage, stage-top incubator with temperature set to 37C and 5% CO2 control, and Andor iXon X3 EMCCD camera with 512 x 512-pixel chip (16-micron pixels). The images were processed and analyzed using imageJ. For immunofluorescence microscopy, cells were fixed using 4% paraformaldehyde for 15 min followed by permeabilization with 0.1% Triton X-100 for 5 min and blocked for an hour in 5% normal goat serum+2% BSA, all in 1XPBS. Primary antibodies were added in 1:200 dilution for 2-3 hours at room temperature or overnight at 4°C which is then incubated with Alexa Fluor conjugated secondary antibodies for 1 hour in dark. These coverslips were mounted in ProLong Gold (Invitrogen) for confocal microscopy. These mounted coverslips were left at room temperature for solidification of ProLong Gold, as suggested by the manufacturer, and imaged on the Dragonfly 200 High-speed Spinning disk confocal imaging platform (Andor Technology Ltd.) on a Leica DMi8 microscope stand equipped with iXon EMCCD and sCMOS Zyla camera. Images were taken under 100x/1.4 oil immersion objective by using Fusion Version 2.3.0.36 (Oxford Instruments) software and deconvolved on associated Imaris software simultaneously. Flow cytometry and competition assay: Cells were detached, harvested in assay media and kept in suspension for 60 min followed by treatment with Fd-sheet in absence or presence of Mn 2+ for 30min. In competition assay with the mb (integrin minibinder?), these cells were first incubated with increasing concentration of mb for 15 min followed by Fd-sheet treatment for additional 15 min and fixed with 4% paraformaldehyde in PBS for 15 min. These cells were washed with 1XPBS trice before flow cytometric analysis on BD FACSymphony 3. FACS files were analyzed using FlowJo software.

### Tight Junction Recovery assay

Tight junction recovery assay was performed according to the protocol described in Zhao et al. 2026 BioRxiv in preparation. Briefly, HUVECs or Human Brain Microvasculature Endothelial Cells (HBMECs) were thawed and cultured on 1% gelatin for two passages before being plated on glass coverslips to grow until 90-100% confluent. Cells were treated with an actin inhibitor, 0.25µM Latrunculin-A (LatA, Cayman, 10010630), diluted in growth media for 30 minutes to disassemble tight junctions. After 30 minutes, LatA was washed out and rinsed with 1X PBS before adding F-domain scaffolds at 100 nM diluted in LG DMEM + 10% FBS for 30 minutes. Cells were fixed with 4% PFA for immunofluorescence staining with antibodies to visualize ZO1 (Invitrogen, 1:100) or Claudin-5 (Abcam, 1:100). Phalloidin (1:100, Invitrogen, A12380) was added during secondary stain. Stained cells were mounted on slides using VECTASHIELD plus DAPI. Phalloidin stains were used to find cell colonies with good contact before switching to ZO1/CLDN5 stains to count junctional tight junction assembly. At least 100 cells were counted per coverslip.

### F-domain Sheet Immunostaining

To analyze F-domain sheets binding assay were performed according to protocol described in Zhao et al. 2021. Briefly, Serum-starved cells were treated with sheets at 50nM of binding domain for 10-30 minutes in low glucose DMEM before fixation with 4% PFA. The fixed cells were washed three times at 5 minutes each before blocking for 1 hour with 3% BSA (VWR, 0332-500G) and 0.1% Triton X-100 (Sigma, T9284-500ML) in PBS while on nutation. The cells were then incubated with primary antibody diluted at 1:100 in blocking agent overnight using these dilutions: Tie2 (Cell Signaling, catalog #4224S), FOXO1 (Cell Signaling, 2880), and α5 integrin (Millipore, 1928), active β1 integrin 9EG7 (BD Biosciences, 550531), inactive β1 integrin mAb13 (BD Biosciences, 552828), phospho-CAS, CAS (Cell Signaling, 4011S), CD144 (Cell Signaling, 2500S), ZO1 (Invitrogen, 33-9100 or 61-7300), Claudin-5 (abcam, ab15106), and Occludin (Invitrogen, 710192).

After the primary antibody, the cells were washed three times at 5 minutes each with 1X PBS while on nutation. The cells were then incubated with secondary antibodies at 1:200 (1:100 for Tie2) diluted in blocking agent for 1 hour and 20 minutes at 37° C. Secondary antibodies were then removed, and cells were washed for three times, 10 minutes each with PBS on nutation. Coverslips were sealed using VECTASHIELD plus DAPI (Vector laboratories, H-2000-2) upside-down on glass slides for analysis in confocal (Leica SP6 or SP8).

### Flow Cytometry analysis of F-domain sheet binding in MCF10A cells

MCF10A were trypsinized and resuspended in assay media. F-domain sheets were added to the cells for 30 minutes and incubated at 37 degrees C with nutation. After incubations, cells were spinned (1600 rpm) and washed once with PBS before fixation with 4% PFA for 15 minutes. Fixed cells were then washed 3 times with 1XPBS, 5 minutes each on nutation. Fixed cells were then transferred to flow cytometry tubes for FACS analysis to detect GFP signal using the FITC detector.

### FOXO Immunostaining

FOXO immunostaining was performed according to protocol described in Zhao et al., 2021. Briefly, serum-starved HUVECs were treated with Tie2 binders for 30 minutes in low glucose DMEM before fixation with 4% PFA. The fixed cells were washed three times at 5 minutes each before blocking for 1 hour with 3% BSA (VWR, 0332-500G) and 0.1% Triton X-100 (Sigma, T9284-500ML) in PBS. After blocking, primary antibody FOXO1 (Cell Signaling, 2880) diluted in blocking agent overnight at 1:100 ratio. After the primary antibody, the cells were washed three times at 5 minutes each with 1X PBS. The cells were then incubated with secondary antibodies at 1:200 diluted in blocking agent for 1 hour and 20 minutes at 37° C. After that, cells were washed for three times at 10 minutes each with PBS. Coverslips were sealed using VECTASHIELD plus DAPI (Vector laboratories, H-2000-2) upside-down on glass slides for analysis in confocal using microscopy Leica SP6 or SP8.

### Endosomal and junctional Tie2 Immunostaining

HUVECs and TIE2-GFP–overexpressing HUVECs were cultured to confluence on glass coverslips coated with either 1 µg/mL fibronectin or 0.1% gelatin for 48 hours. Confluent monolayers were serum-starved in conditioned medium diluted 1:10 in basal endothelial growth medium (without supplements) for 3 hours. Cells were then treated with 10 nM or 20 nM of H8F, H8T, or Comp-ANG1 (used as a control) in starvation medium for 15–30 minutes. Following treatment, cells were fixed with 4% paraformaldehyde (PFA) in PBS for 10 minutes at room temperature. For surface staining, non-permeabilized HUVECs were labeled for surface TIE2 and inactive β1-integrin (mAb13). After surface staining, cells were permeabilized with 0.3% Triton X-100 in PBS and subsequently stained for FOXO1 and VE-cadherin (CDH5). For intracellular staining, TIE2-GFP–overexpressing HUVECs were permeabilized with 0.3% Triton X-100 in PBS after fixation and stained for phospho-TIE2 and EEA1.

Coverslips were imaged using a Leica Stellaris 8 confocal microscope equipped with a 63× oil immersion objective, or Leica Stellaris 5 confocal microscope equipped with a 63× glycerol immersion objective. Images were acquired in Lightning mode with a zoom factor of 1.4 and a resolution of 2048 × 2048 pixels.

All image analyses were performed using ImageJ (Fiji). To quantify the junction-to-membrane ratio of TIE2 or mAb13, a binary mask of endothelial junctions was generated based on CDH5 immunostaining. The integrated density of TIE2 or mAb13 signal within this junctional mask was measured and defined as the junctional signal. Signal located outside the junctional mask was considered membrane-associated. The junction-to-membrane ratio was calculated by dividing the junctional signal by the membrane signal.

To determine the nuclear-to-cytoplasmic ratio of FOXO1, a binary nuclear mask was created based on DAPI staining. The integrated density of FOXO1 signal within the nuclear mask was measured and defined as the nuclear signal, while the signal outside the nuclear mask was considered cytoplasmic. The nuclear-to-cytoplasmic ratio was calculated by dividing the nuclear signal by the cytoplasmic signal.

Internalized TIE2-GFP/EEA1 and phospho-TIE2/EEA1 vesicles were analyzed using ImageJ with an object-based colocalization approach. Each channel was segmented and converted into binary masks, which were then combined using the Process > Math > AND function to identify overlapping pixels. Colocalizing vesicles were quantified using the Analyze Particles tool and normalized to the total cell count (nuclei). Statistical significance was assessed using ordinary one-way ANOVA followed by post hoc pairwise comparisons with Tukey’s multiple comparison test.

### Tie2 Minibinder in vivo administration and influenza challenge in mice

Animal studies were approved by the University of Washington Institutional Animal Care and Use Committee under protocol number 4266-01. Female, 6–10 week-old BALB/c mice were randomly assigned into groups, anesthetized, and 30 ul of saline or the H8mb was intranasally (IN) administered at 5.0 mg/kg at one hour post-influenza infection and then repeated for 4 subsequent administrations at the same concentration every 24 hours. Mice were anesthetized with 2.5-4.5% isoflurane and challenged 30 ul IN with a 100xTCID50 dose of A/California/04/09 (H1N1) as described (PMID: 26845438). The mice were monitored daily for weight loss and survival for a maximum of 14 days post-infection. Animals that lost more than 30% of their initial body weight were euthanized by carbon dioxide in accordance with our animal protocols. At least five mice per group were used for each experiment. All mice used for the experiments are included for analyses. For mouse experiments, researchers were not blinded to animal identity.

#### Acknowledgments

We are grateful to the staff of the Research Solution Center at IBS for help with cryo-EM data collection. Cryo-EM data processing was performed on Olaf, the data analysis hub in the IBS Research Solution Center. We are grateful to Biomedicum Imaging Unit, Helsinki Institute of Life Science (HiLIFE) for microscopy services. We acknowledge Howard Hughes Medical Institute (DB) for their support.

## Funding

National Institutes of Health grant R01CA114536 (CM)

National Institute of Dental and Craniofacial Research T90DE021984 (YTZ)

National Institutes of Health DE033016 (HRB)

National Institutes of Health 1P01GM081619 (HRB)

National Institutes of Health R01GM097372 (HRB)

National Institutes of Health R01GM083867 (HRB)

National Institutes of Health R01AI160052 (HRB)

National Heart, Lung, and Blood Institute Progenitor Cell Biology Consortium U01HL099997 (HRB)

National Heart, Lung, and Blood Institute Progenitor Cell Biology Consortium UO1HL099993 (HRB)

National Institutes of Health Somatic Cell Genome Editing COF220919 (HRB)

Stem Cell Gift Fund (HRB)

Department of Defense PR203328 W81XWH-21-1-0006 (DB, HRB)

National Research Foundation of Korea (RS-2024-00397681 (HMK)

InnoCORE program of the Ministry of Science and ICT N10250153 (HMK)

KAIST Convergence Research Institute Operation Program (YF)

Wihuri Foundation (PS)

Sigrid Jusélius Foundation (PS)

Academy of Finland Centre of Excellence Program 346134 (PS)

## Author contributions

Conceptualization: CM, YTZ, DB, HRB

Minibinder design & screening: BC, IG, CM, YTZ

Cryo-EM: FY

Experimental Strategies: CM, YTZ, SK, AP, PB, SZ

Visualization: CM, YTZ, SK, FY, AP

Funding acquisition: DB, HRB

Supervision: DB, HRB

Writing – original draft: CM, YTZ, HRB

Writing – review & editing: All

## Competing interests

A provisional patent covering Tmb1 and all minibinder sequences presented in this paper has been filed by the University of Washington. All other authors declare no competing interests.

## Data, code, and materials availability

All data are available in the main text or the supplementary materials.

## Supplementary Materials

Figs. S1 to S9

Tables S1 to S3

## Notes

### Competing Interest Statement

The authors have declared no competing interest.

