## Supplemental Figures for "Integrin-independent Tie2 activation using de novo designed proteins"

**Supplementary Figures S1-S9**

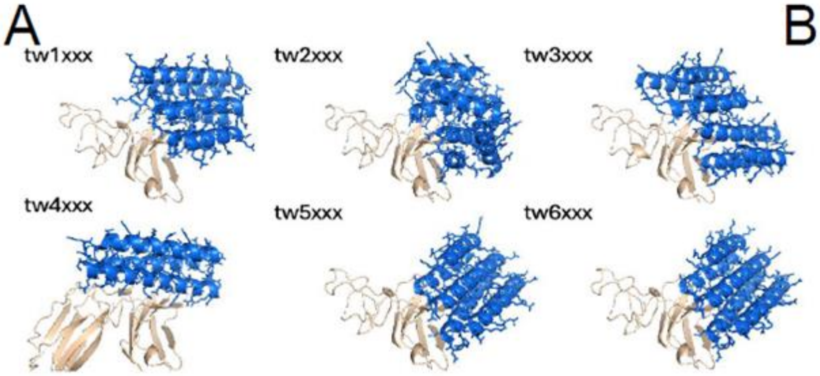


B

| | Name | KD (nm) |
|---|---|---|
| Family 1 | Tie2_tw1102 | 0.295 |
| | Tie2_tw1202 | 0.853 |
| | Tie2_tw1301 | 0.615 |
| Family 2 | Tie2_tw2101 | 2.09 |
| | Tie2_tw2106 | 2 |
| | Tie2_tw2201 | 1.01 |
| | Tie2_tw2202 | 0.258 |
| Family 4 | Tie2_tw4000 | 28.3 |
| | Tie2_tw4104 | 6 |
| Family 5 | Tie2_tw5104 | 0.785 |
| Family 6 | Tie2_tw6103 | 3.61 |
| | Tie2_tw6105 | 1.19 |
| | Tie2_tw6107 | 4.18 |

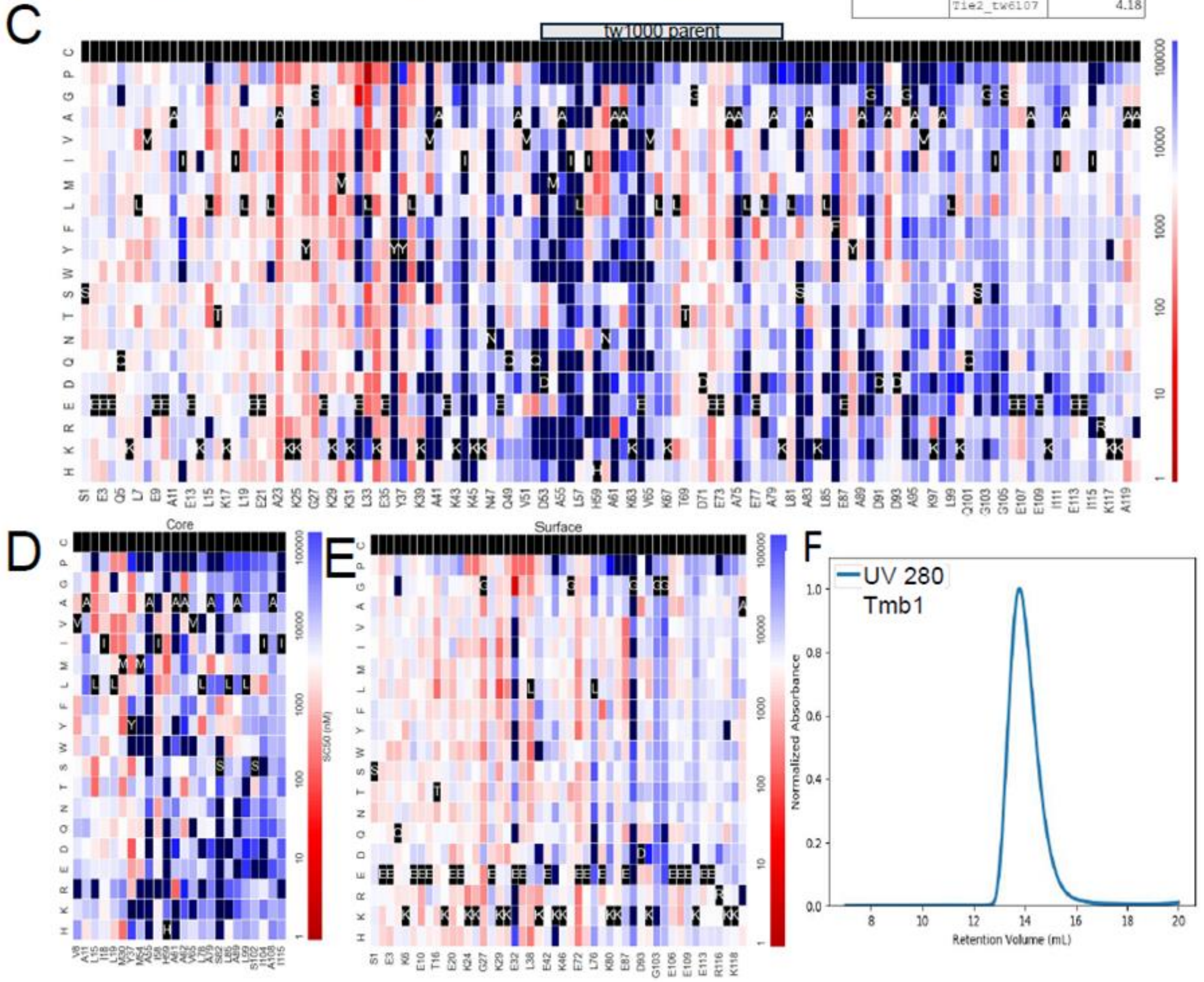


**Figure S1 Tie2 mini binder design variants.**
A) The 6 families of Tie2 binders. Shown are the original twX000 binders that are the parents of each families in complex with Tie2. Each family is derived from these binders via a series of mutations. Secondary Structure Matching (SSM) graphs of tie2 mini binder designs. C) The full SSM graph of tw1000 with red positions showing better affinity than the parent tw1000 design, affinity as quantified by SC50 (the concentration where 50% of the expressing cells are collected), color scale is from red to blue indicating affinity from high to low respectively. D). The subset of positions at the interface core. This graph shows less conservation than one might initially expect for a protein binder, but the larger proteins tend to show less core conservation in

our experience as there is so much free energy available for folding that a single mutations cannot disrupt the protein. The core also shows many mutations with room for improvement which is likely as a result of the polar nature of the interface. Small backbone changes lead to improved hydrogen bonds at the interface. E) The subset of positions on the protein surface. As expected, the vast majority of surface positions for tw1000 do not show a strong preference in amino acid. Certainly a few matter, but compared to the interface and core, there is much less conservation giving strong suggestion of the fold of the protein. F) SEC trace of the monomeric Tmb1 showing a single monodisperse peak in the UV280 wavelength.

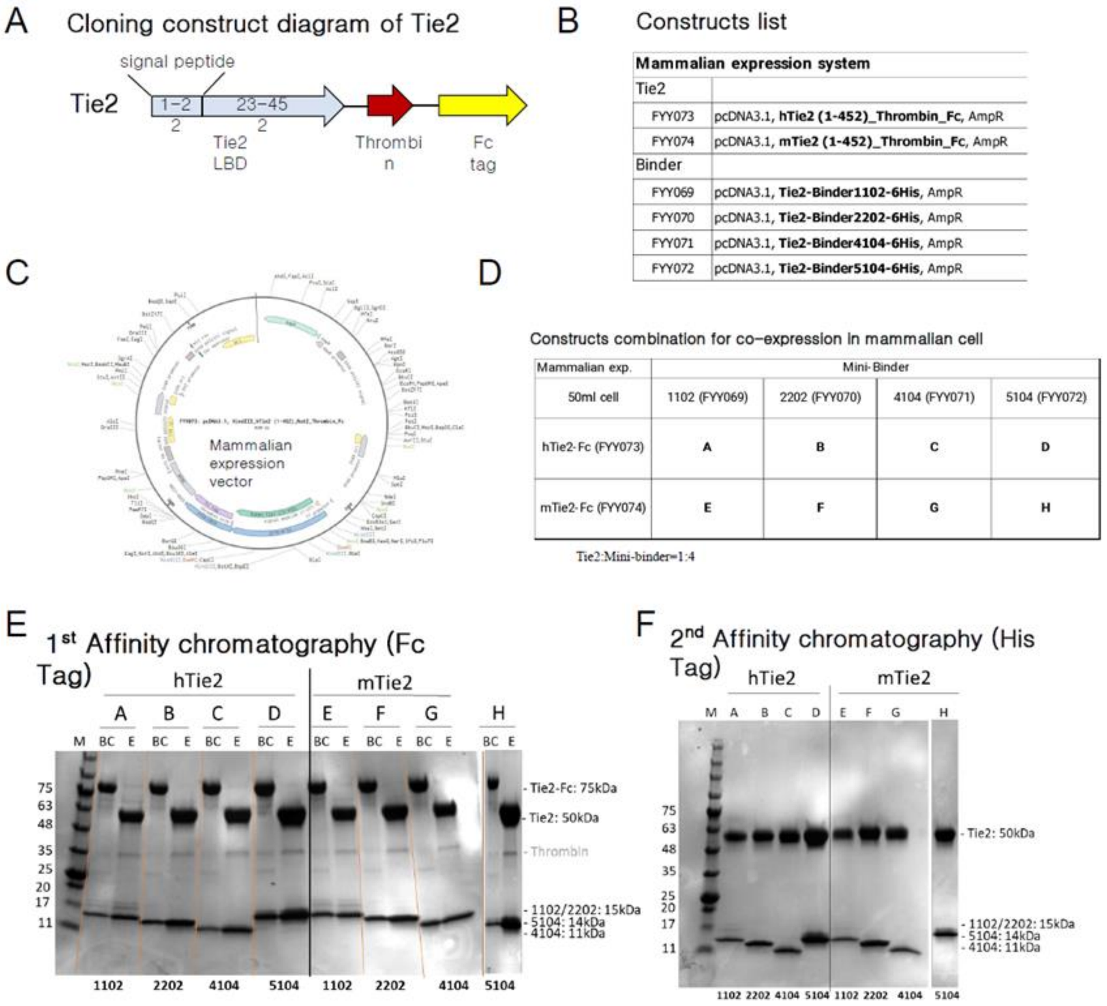


**Figure S2 Expression of minibinder for cryo-EM studies**
A) Cloning construct diagram of the Tie2 ligand binding domains expressed for co-purification. Two versions were made - one with human Tie2 and the other with mouse Tie2. B) Constructs produced for co-purification experiments. C) Mammalian expression vector map for the hTie2 LBD showing the receptor sequence, thrombin site, and fc-tag. A similar construct was made for the mouse-Tie2 LBD. D) Table describing the constructs co-expressed. For example, Tmb1 (1102) was expressed with both hTie2-Fc (and subsequently labeled as A) and mTie2-Fc (labeled as E). E) SDS-PAGE gel of binders co-expressed with hTie2 or mTie2. Letters (A, B, C, etc) represent the binder-receptor pairings defined in D. Lanes are labeled as BC for Before Column and E for elution to indicate whether the same had been passed over a Protein A column. (See: methods). F) SDS-PAGE gel of constructs after nickel column elution. Binder and receptor LBD eluted together, indicative of binding between the two.

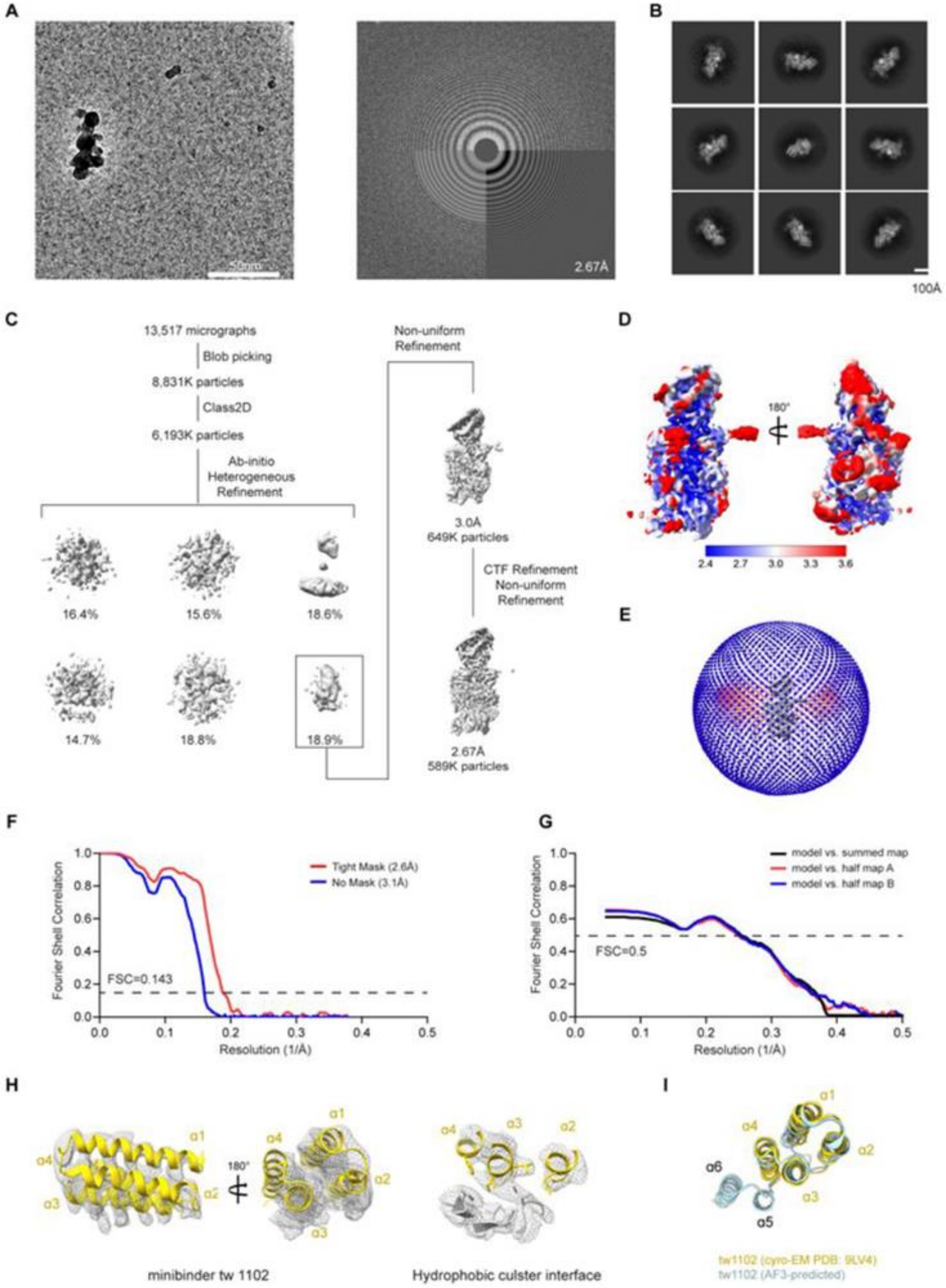


**Fig. S3 Cryo-EM analysis of 4 helix Tmb1**

(A) Representative cryo-EM micrograph (left) and its Fourier transform (right) of the hTie2 LBD-Tmb1 complex. (B) Representative 2D class averages of the hTie2 LBD-Tmb1 complex. (C) Data processing workflow of hTie2 LBD-Tmb1 complex. (D) Final cryo-EM map colored with local resolution. (E) Euler angle distribution of all particles in the final 3D reconstructions. The cylinder bars' height and color (from blue to red) are proportional to the number of particles in those views. (F) Gold-standard Fourier shell correlation (FSC) for two independently refined half maps in cryoSPARC (resolution cutoff at FSC=0.143). (G) FSC curves for the refined model of hTie2-Tmb1 complex versus the map (resolution cutoff at FSC = 0.5). (H) Cryo-EM map density and model of hTie2-Tmb1 complex. (I) Comparison of experimental and AlphaFold3-predicted structures of the minibinder Tmb1. The Cα RMSD for Tmb1 only is 0.8 Å.

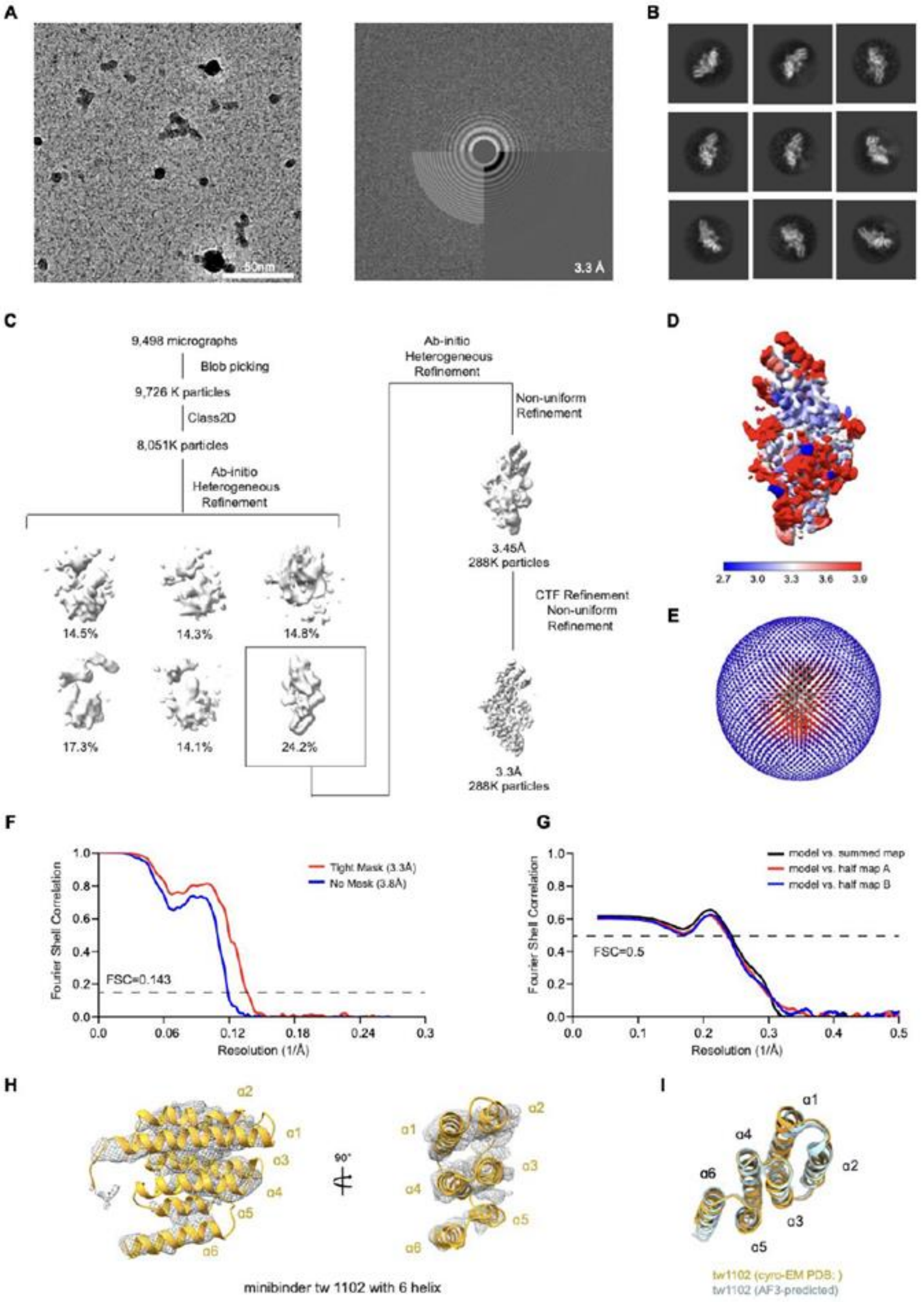
A
50nm
3.3 Å
B
C
9,498 micrographs
Blob picking
9,726 K particles
Class2D
8,051K particles
Ab-initio
Heterogeneous
Refinement
14.5%
14.3%
14.8%
17.3%
14.1%
24.2%
Ab-initio
Heterogeneous
Refinement
Non-uniform
Refinement
3.45Å
288K particles
CTF Refinement
Non-uniform
Refinement
3.3Å
288K particles
D
2.7 3.0 3.3 3.6 3.9
E
F
Tight Mask (3.3Å)
No Mask (3.8Å)
FSC=0.143
Fourier Shell Correlation
Resolution (1/Å)
G
model vs. summed map
model vs. half map A
model vs. half map B
FSC=0.5
Fourier Shell Correlation
Resolution (1/Å)
H
α1
α2
α3
α4
α5
α6
90°
minibinder tw 1102 with 6 helix
I
tw1102 (cyro-EM PDB: )
tw1102 (AF3-predicted)

**Fig. S4 Cryo-EM analysis of 6 helix Tmb1**
(A) Representative cryo-EM micrograph (left) and its Fourier transform (right) of the hTie2 LBD-minibinder Tmb1 six-helix complex. Tmb1 is also known as 1102. (B) Representative 2D class averages of the hTie2 LBD-minibinder Tmb1 complex. (C) Data processing workflow of hTie2 LBD-Tmb1 complex. (D) Final cryo-EM map colored with local resolution. (E) Euler angle distribution of all particles in the final 3D reconstructions. The cylinder bars' height and color (from blue to red) are proportional to the number of particles in those views. (F) Gold-standard Fourier shell correlation (FSC) for two independently refined half maps in cryoSPARC (resolution cutoff at FSC=0.143). (G) FSC curves for the refined model of hTie2-minibinder Tmb1 complex versus the map (resolution cutoff at FSC = 0.5). (H) Cryo-EM map density and model of hTie2-Tmb1 complex. (I) Comparison of experimental and AlphaFold3-predicted structures of the minibinder Tmb1.

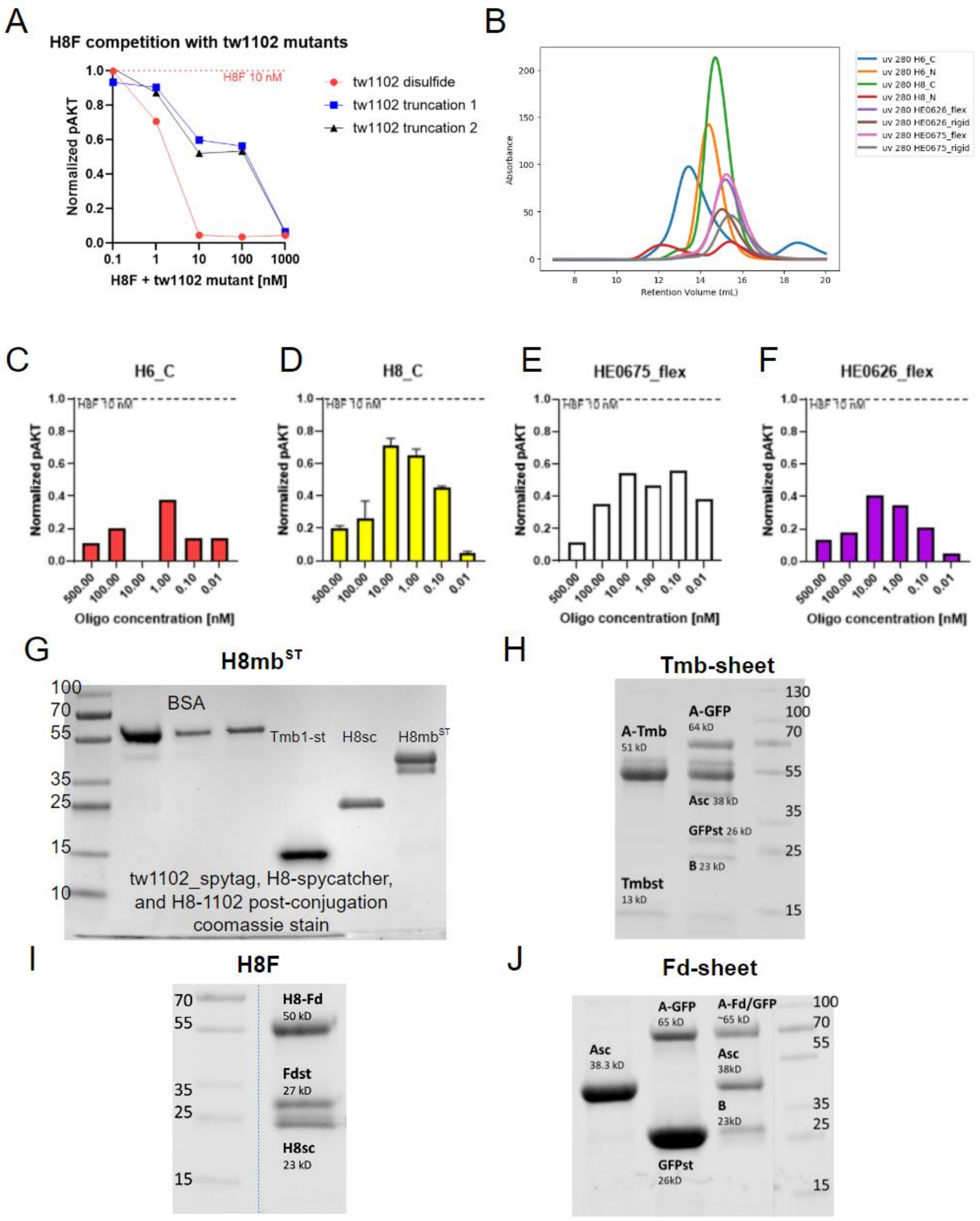


**Figure S5 Tmb1-ST conjugation to scaffolds and screening of alternate oligomeric**

**assemblies**

A) Truncation mutants of tw1102 are not as good at outcompeting F-domain mediated activation of Tie2, indicating a reduced affinity to the Tie2 ligand binding domain. B) SEC traces of oligomerized Tie2 minibinder molecules. C-F) Activity of selected Tie2 oligomers. G-J) Coomassie blue staining of Tmb-st or F-domain-st conjugated to H8sc scaffold or Asc domain of sheets using SpyCatcher (sc)-SpyTag(st). GFP-st is conjugated to Asc for Tmb/Fd-sheet binding analysis in confocal imaging and flow cytometry. Conjugations that achieved >90% efficiency were used for all experiments.

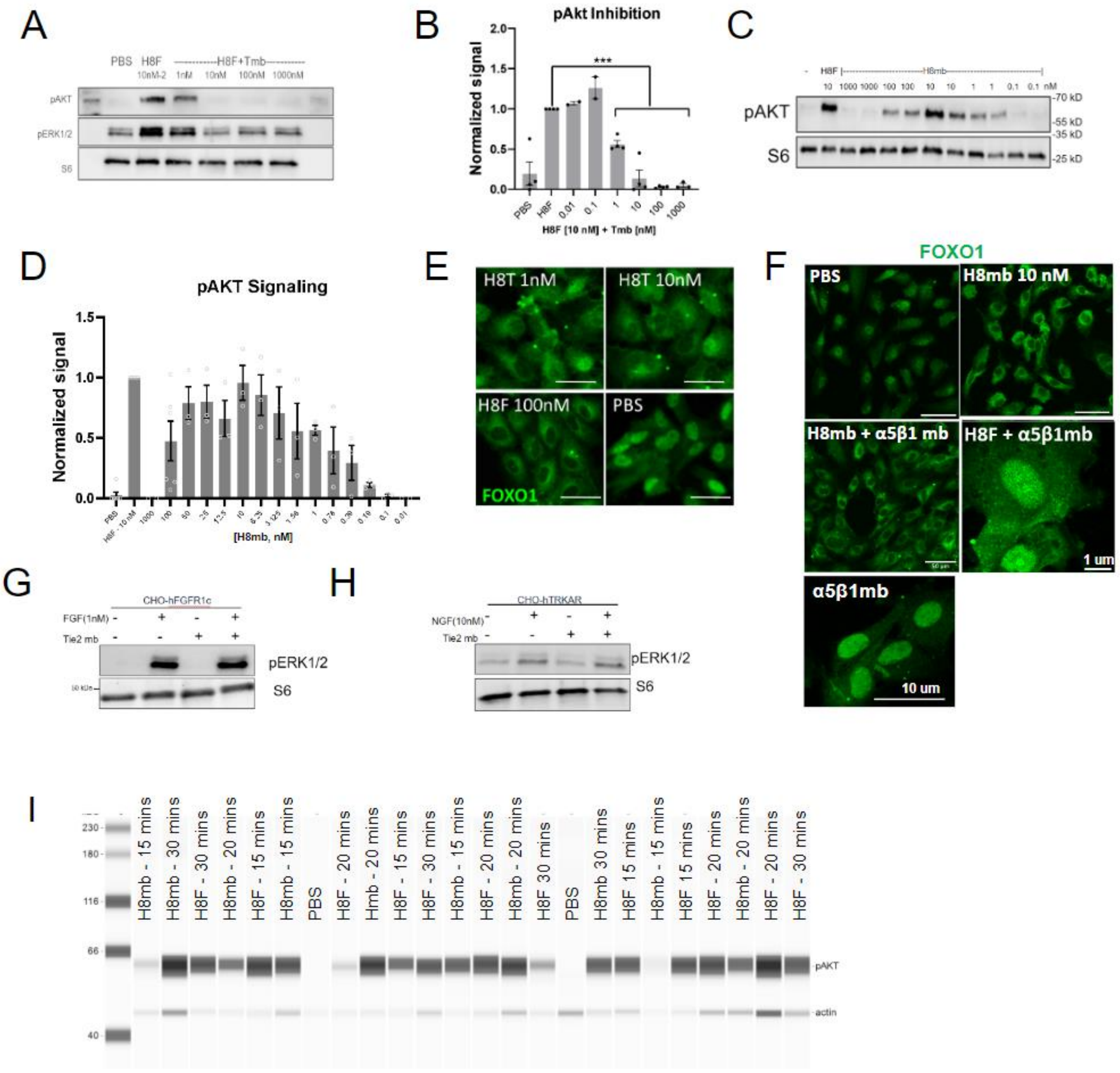


**Figure S6 Characterization of Tmb1, H8mb, and representative images and western blots**
A) Representative western blot staining of pAKT and pERK signaling in competition experiment between H8F at 10 nM +/- Tmb from 1 to 1000 nM in serum starved HUVECs. B) Quantification of pAKT/S6 level in competition experiments normalized to 10 nM of H8F samples as positive control. C) Competition data in B were replotted to estimate IC50 using GraphPad Prism. D) Representative images of pAKT activation upon 10 nM of H8F, H8mb or H3T administration in serum starved HUVECs. E) Quantification of pAKT/S6 level upon H8mb treatment normalized to H8F samples, selected subsets of this data were re-plotted in Fig. 3C to estimate $EC_{50}$. E-F) Immunofluorescence images of FOXO1 staining in HUVECs, size bars are 10um. G-H) Western blot staining showing Tmb does not interact with other RTKs, thus Tmb was not able to inhibit FGF or NGF ligand activity in their respective receptor overexpression CHO lines. I) Representative pseudo-blot of the time course H8mb and H8F experiment. Ladder in kDa on

the left and samples contain labeled bands for actin and pAKT.

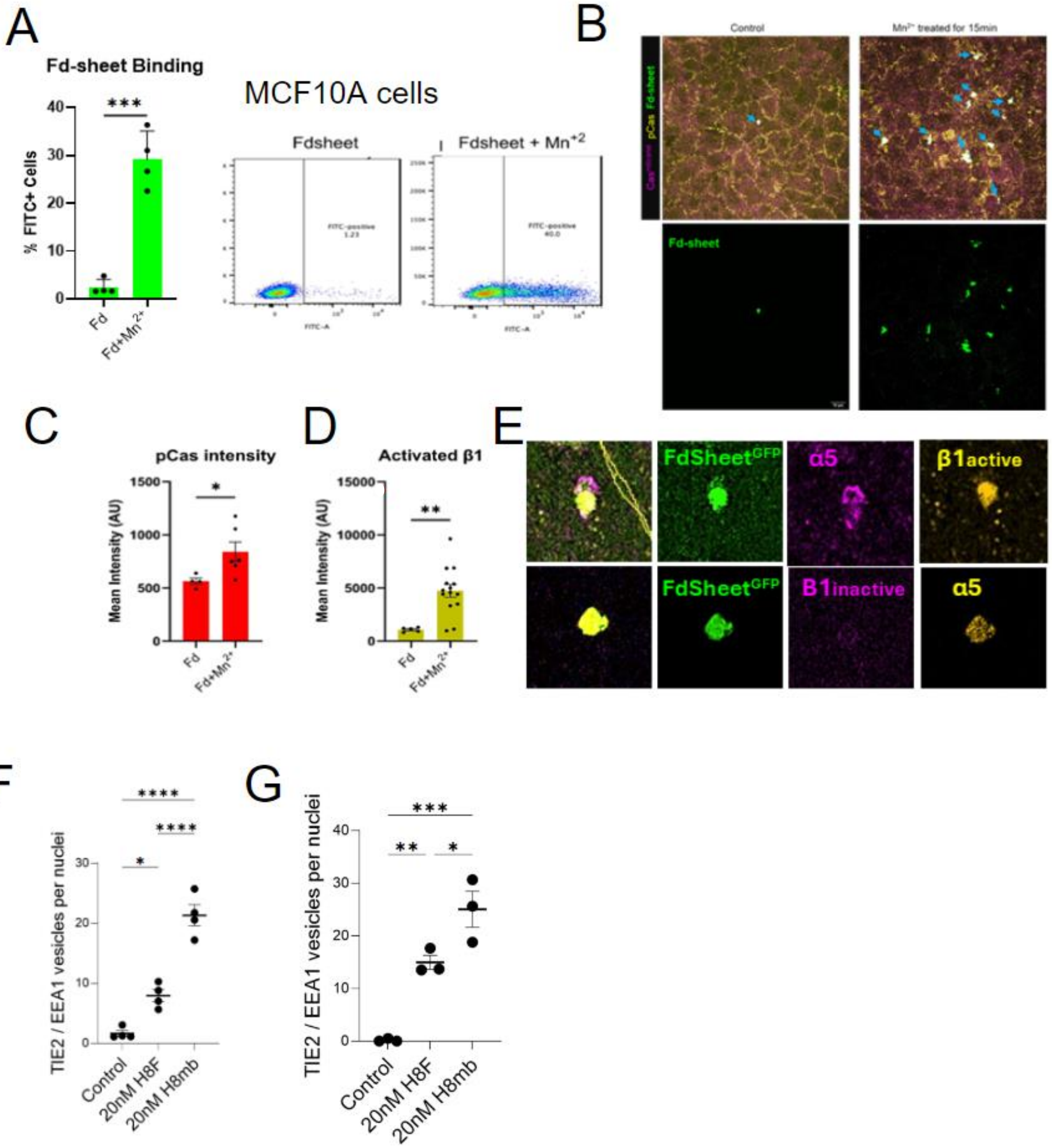


**Figure S7 - F-domain bids MCF10A cells and recruits integrin complex components**
A) Flow cytometry analysis of MCF10A cells incubated with GFP-tagged F-domain sheets. FITC intensity on x-axis, side scatter on y-axis. B) Representative images of MCF10A cells incubated with GFP tagged F-domain sheets with or without $Mn^{2+}$. Top row: merged images, bottom row: GFP channel only. C) Mean intensity for pCas in Fd treated cells with and without $Mn^{2+}$. Each dot represents one biological replicate. D) Mean intensity for activated beta1 in Fd treated cells with and without $Mn^{2+}$. Each dot represents one biological replicate. E) Representative immunofluorescence of MCF10A cells quantified in Figure 3E. F-G) Quantification of Tie2-GFP co-localized with EEA1 signal from images represented in Figure 4 at 15 minutes (left) and 30 minutes (right).

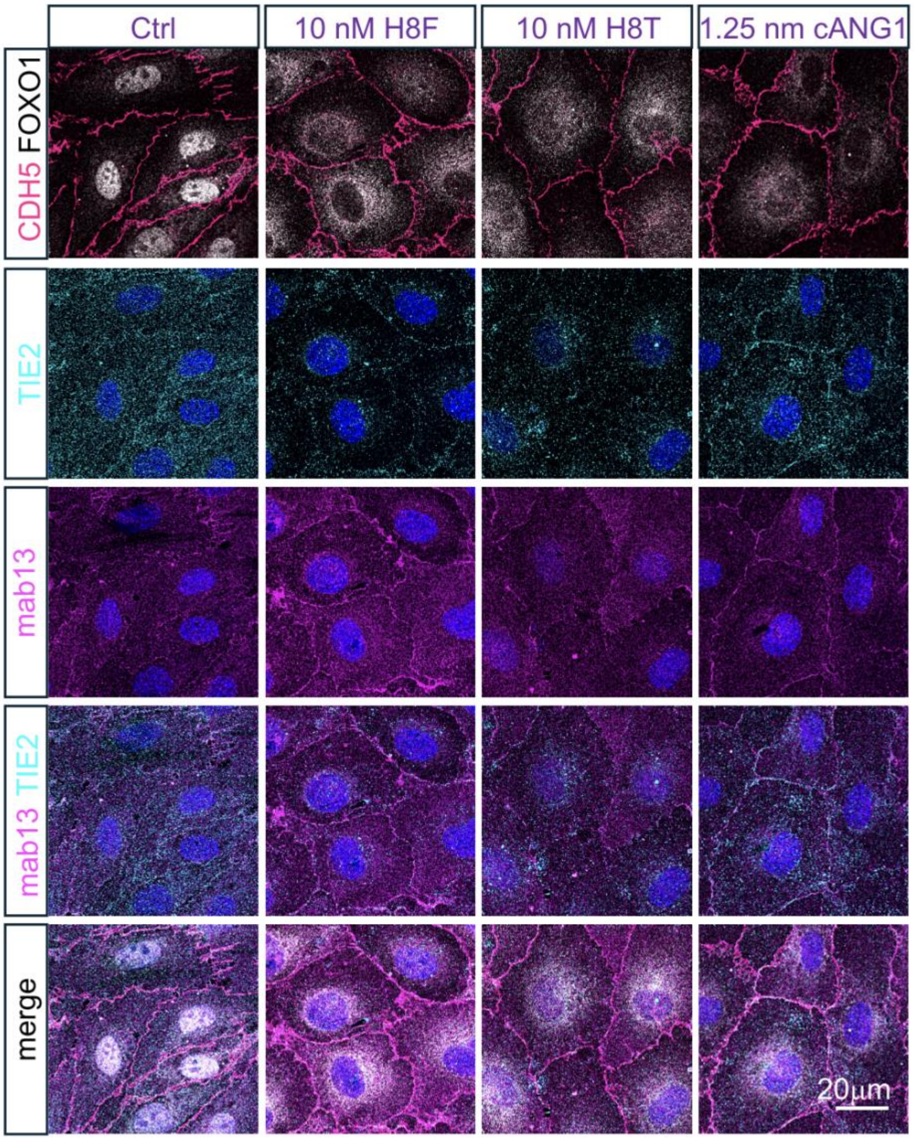


**Figure S8 - Lack of endogenous Tie2 and beta1-integrin in cell-cell junctions after H8mb stimulation (30 minutes).** Representative images of confluent HUVECs fixed after 30 minute incubation with PBS (Ctrl), 10 nM H8F, 10 nM H8mb, or 1.25 nM cANG1. Top row: CDH5 (pink) staining used to create a binary mask (see methods). FOXO1 localization in white. 2nd row: endogenous Tie2 stain. 3rd row: mab13 staining which is specific to the inactive form of beta1 integrin. 3rd row: Merged images of Tie2 and mab13. Bottom row: Merge of all channels. Quantitative analysis found in Figure 4F-H. Scale bar represents 20 μm.

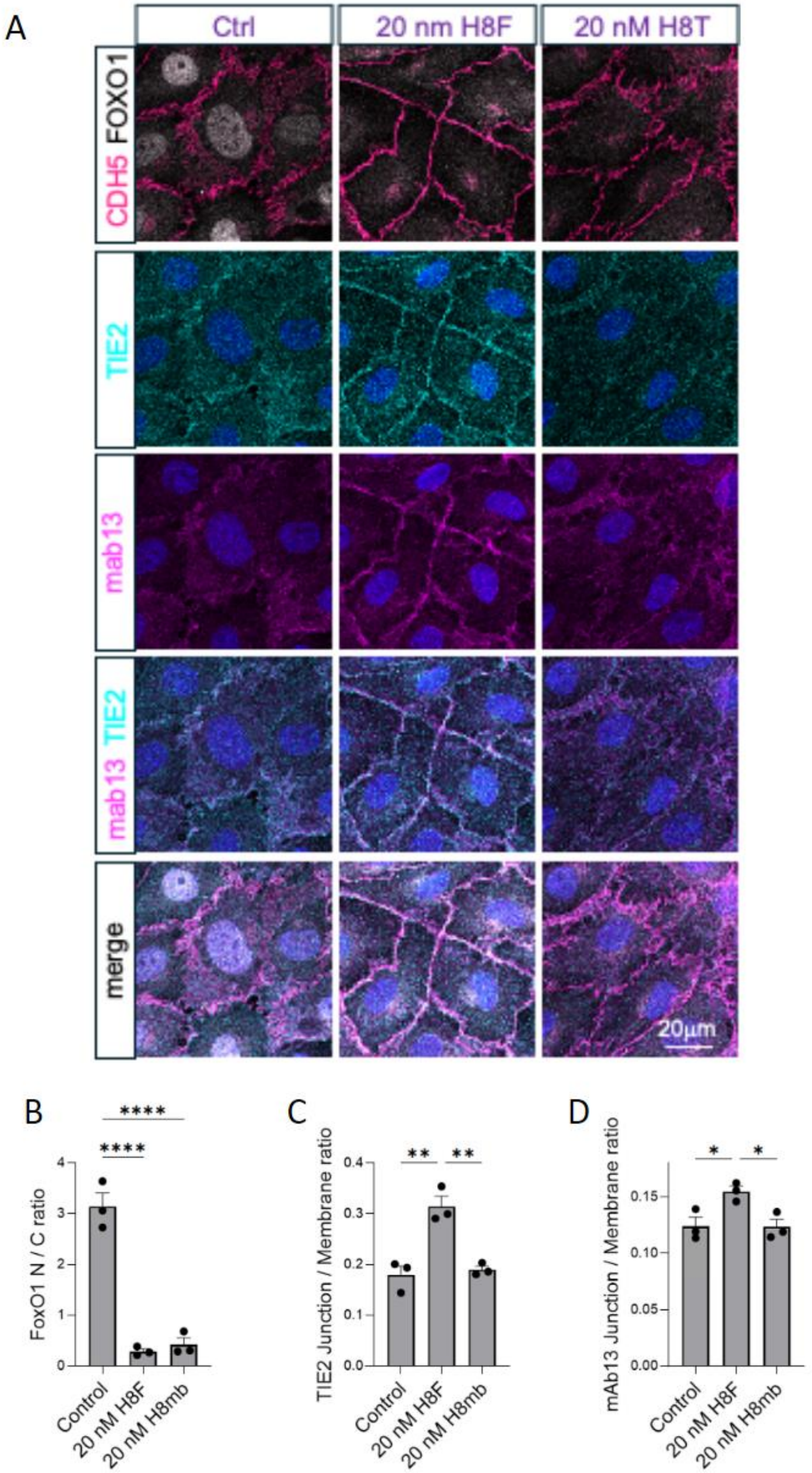
A
Ctrl
20 nm H8F
20 nM H8T
CDH5 FOXO1
TIE2
mab13
mab13 TIE2
merge
20μm
B
FoxO1 N / C ratio
Control
20 nM H8F
20 nM H8mb
C
TIE2 Junction / Membrane ratio
D
mAb13 Junction / Membrane ratio

**Figure S9 - Lack of endogenous Tie2 and b1-integrin in cell-cell junctions after H8mb stimulation (15 minutes)**

A) Representative images of confluent HUVECs fixed after 15 minute incubation with (from left to right) PBS (Ctrl), 20 nM H8F, or 20 nM H8mb. Top row: CDH5 (pink) staining used to create a binary mask for junctional/non-junctional analysis (see methods). FOXO1 localization in white. 2nd row: endogenous Tie2 stain. 3rd row: mab13 staining which is specific to the inactive form of beta1 integrin. 3rd row: Merged images of Tie2 and mab13. Bottom row: Merge of all channels. Scale bar represents 20 µm. B) Ratio of nuclear to cytoplasmic FOXO1 signal intensities across three experiments. C) Quantitative analysis of Tie2 signal intensity reported as a ratio of junctional vs non-junctional signal. D) Quantitative analysis of inactive β1 integrin location reported as a ratio of junctional vs non-junctional signal. All statistical significances were assessed using ordinary one-way ANOVA followed by post hoc pairwise comparisons with Tukey's multiple comparison test.

| Designs | Sequences |
|---|---|
| >Tie2_tw1000 | SEEEQKLVEEAIEKLTKILEELAKKYGEKMKELKEYYLKVAEKIKKNEQAVQDMAILIH<br>NAAKEVLKLTGDEEAALELAKLSAKLFEYAGDADGAVKALKQSGIGEEAEKIAEEIRK<br>KAA |
| >Tmb_1 | SEEEQKLVEEAIEKLTKILEELAKKYGEKMKEPKEYYLRMSEKIKKNEQPEQDMAILIH<br>NAGKEVLKLTGDEEAALELAKLSAKLFQYAGDTDGAVRALKQSGIGEEAEKIAEEIRK<br>KAA |
| >Tie2_tw1202 | SEEEQKLVEEAIEKLTKILEELAKKYGEKMKEPKEYYLRMAEKIKKNEQPEEDMAILIH<br>NAGKEVLKVTGDEEAALELAKLSAKLFQYAGDTAGAVRALKQSGIGEEAEKIAEEIRK<br>KAA |
| >Tie2_tw1301 | SEEEQKLVEEAIEKLTKILEELAKKYGEKMKEPKEYYHRMSEKIKKNEQAEQDMAILL<br>HNACKEVMKLTGDEEAALELAKLSAKLFEYAGDTDGAVRALKQSGIGEEAEKIAEEIR<br>KKAA |
| >Tie2_tw2000 | DRLREIIELAREAAEEGLSPAVAALAARRATGDDVAAIIANLMAHAGGENAERVARV<br>VWETSERLGGSSVRHQDIALAVGRAVLYRLRGDEEEAEFYEKLALKIAKREEDRKLV<br>KEILEE |
| >Tie2_tw2101 | DRLREIIELAREAAEEGQSPEEASRRARRATGNGVAAIIVHLMAHAGGENAERVARVV<br>WETSERLGGSSEKHQDIALTVGRAVLYRLRGDEEEAEYYEKLALKIAKREEDRKLVK<br>EILEE |
| >Tie2_tw2106 | DRLREIIELAREAAEEGLSPELASRRARRATGDGVAAIIVHLMAHAGGENAERVARVV<br>WETSERLGGSSERHQDIALTVGRAVLYRLRGDEEEAEYYEKLALKIAKREEDRKLVKE<br>ILEE |
| >Tie2_tw2202 | DRLREIIELAREAAEEGLSPAEAALRARRATGNGVAAIIAHLMAHAGGENAERVARVV<br>WETSERLGGSSERHQDIALTVGRAVLYRLRGDEEEAEYYEKLALKIAKREEDRKLVKE<br>ILEE |
| >Tie2_tw3000 | NSEKVLKMAEEVAKKTGSETAKKVLENIKEDIKNGEDLTLSAIDAALLLSKIDPEKAV<br>EFLRELGLDRDADVLEVYVYGRKLYEETGSEEIRKAAEAETLAVALGRITAEEALKRL<br>EELV |
| >Tie2_tw4000 | SEEEKERGEKMVEQYAENLRKLAEEYIERGEPPEEILRRIEKEAEAYLEDLETIFEGSEL<br>KEEILELAEKKFEEVKEEVEERL |
| >Tie2_tw4104 | SEEEKERGEKMVEQYSENLRKLAEENIDRGEPPEEIWRRIEKEAEAFLEDLETFFEGRE<br>LKEEILELAEKKFEEVKEEVEERL |
| >Tie2_tw5000 | GVEEPIRELEEYGRRRIERVEKALPGNRLAVLAMRSLAEHLIARVRYAAERGKDVEGL<br>VEAAKLALDVAEAAILKNGLVPGDVADKVLLAFREAEEDPKHAKERLEELKEEV |
| >Tie2_tw5104 | GVEEPIRELEEYGRRRIERVEKALPGNRLAVLSTRSLAKHFISRVRYDAERGKDVEGLV<br>EAAKLALDVADAAIHKNGLVPGDVADKVLLAFREAEEDPKHAKERLEELKEEV |
| >Tie2_tw6000 | GSEELIEELERYGKEVIERIKKALPGNRLAVLAMESFVEHLLKLLEYAAEKGLDVEGIT<br>KVAKLGLEVFEKAILKNGSVPGDIADKVLLAVREAEKDPEEALKRLEEILKEV |
| >Tie2_tw6103 | GSEELIEELERYGKEVIERIKKALPGNRMGVLAMESFVKHLLKLLEYAAEKGLDVEGIT<br>KVDKLGLEVFEKAILKNGSMPGDIADKVLLAMREAEKDPEEALKRLEEILKEV |
| >Tie2_tw6105 | GSEELIEELERYGKEVIERIKKALPGNRLGVLAMQSFVKHLRKSLKYAAEKGLDVEGIT<br>KVAKLGLEVFEKAIHKNGSVPGDIADKVLLAVREAEKDPEEALKRLEEILKEV |
| >Tie2_tw6107 | GSEELIEELERYGKEVIERIKKALPGNRMAVLSMQSFVEHLRKSLEYAAEKGLDVEGIT<br>KVDKLGLEVFEKANHKHGSVPGDIADKVLLAVREAEKDPEEALKRLEEILKEV |

**Supplemental Table 1: Amino acid sequence of Tie2 minibinders.**

Tie2_tw1XXX-6XXX are parent designs. Each parent design has its own family of minibinders derived from the results of an SSM library (see methods). Tmb1 is a child design of tw1000.

| | **hTie2-minibinder Tmb1** |
|---|---|
| **Data collection** | |
| Microscope | TFS Krios G4 |
| Voltage (kV) | 300 |
| Detector | Gatan K3 |
| Magnification | 130,000 |
| Electron dose ($e^{-}/Å^{2}$) | 62.0 |
| Defocus range (μM) | -0.7 ~ -1.8 |
| Pixel size (Å) | 0.648 |
| Micrographs (no.) | 13,517 |
| **Data processing** | |
| Initial particle images (no.) | 1,206,082 |
| Final particle image (no.) | 589,066 |
| Map resolution (Å) | 2.67 |
| FSC threshold | 0.143 |
| **Refinement** | |
| Initial model used (PDB ID) | 2GY5 |
| Model resolution (Å) | 2.7 |
| FSC threshold | 0.143 |
| Model composition | |
| Protein residues | 525 |
| Ligands | 0 |
| B factors ($Å^{2}$) | |
| Protein | 123.14 |
| Ligand | 0 |
| R.m.s deviations | |
| Bond lengths (Å) | 0.005 |
| Bond angles (°) | 0.924 |
| **Validation** | |
| MolProbity score | 2.4 |
| Clash score | 23.5 |
| Poor rotamers (%) | 0 |
| Ramachandran plot | |
| Favored (%) | 90.4 |
| Allowed (%) | 9.6 |

| | |
|---|---|
| Disallowed (%) | 0 |
| **Accession codes** | |
| EMBD code | 63412 |
| PDB code | 9LV4 |

**Supplemental Table 2. Cryo-EM data collection, refinement, and validation statistics**

| Minibinder Tmb1 | | hTie2 | Type | Distance (Å) |
|---|---|---|---|---|
| Helix α2 | N50 | G196-O | Hydrogen bond | 2.8 |
| Helix α3 | D56 | R192 | Salt bridge | 3.5 |
| | | Y156 | Hydrogen bond | 2.6 |
| | N63 | S164 | Hydrogen bond | 2.6 |
| | | D152 | Hydrogen bond | 2.7 |
| | E67 | R167 | Salt bridge | 2.8 |
| Helix α2 | M43 | I194 | Hydrophobic cluster | 3.7 |
| Helix α3 | L60 | I194 | | 4.2 |
| | I59 | I194 | | 3.5 |
| | | V154 | | 3.5 |

| | | | | |
|---|---|---|---|---|
| | | Y156 | | 3.7 |
| | | F161 | | 3.7 |
| Helix α4 | F89 | F161 | | 3.7 |

**Supplementary Table 3. Interactions of minibinder Tmb1 with Tie2 LBD.** The interactions were identified using PDBe PISA (for salt bridge and hydrogen bond) and BIOVIA Discovery Studio Visualizer (for hydrophobic interaction). The cut-off distances are following the default settings in PDBe PISA, and distance measurements were performed in COOT.
